# AlphaFold2 SLiM screen for LC3-LIR interactions in autophagy

**DOI:** 10.1101/2024.09.06.611604

**Authors:** Jan F. M. Stuke, Gerhard Hummer

## Abstract

In selective autophagy, cargo recruitment is mediated by LC3-interacting regions (LIRs) / Atg8-interacting motifs (AIMs) in the cargo or cargo receptor proteins. The binding of these motifs to LC3/Atg8 proteins at the phagophore membrane is often modulated by post-translational modifications, especially phosphorylation. As a challenge for computational LIR predictions, sequences may contain the short canonical (W/F/Y)XX(L/I/V) motif without being functional. Conversely, LIRs may be formed by non-canonical but functional sequence motifs. AlphaFold2 has proven to be useful for LIR predictions, even if some LIRs are missed and proteins with thousands of residues reach the limits of computational feasibility. We present a fragment-based approach to address these limitations. We find that fragment length and phosphomimetic mutations modulate the interactions predicted by AlphaFold2. Systematic fragment screening for a range of target proteins yields structural models for interactions that AlphaFold2 and AlphaFold3 fail to predict for full-length targets. We provide guidance on fragment choice, sequence tuning, and LC3 isoform effects for optimal LIR screens. Finally, we also test the transferability of this general framework to SUMO-SIM interactions, another type of protein-protein interaction involving short linear motifs (SLiMs).

## Introduction

Selective autophagy is central to homeostasis in eukaryotic cells. Autophagy is used to rid cells of pathogens [1], recycle organelles [2, 3], and degrade protein condensates [4] and aggregates [5]. The cargo is enveloped by a double membrane, the phagophore, and then degraded upon phagophore fusion with a lysosome [6]. Selective autophagy is associated with several pathophysiological processes, including neurodegenerative diseases and cancer [7].

LC3 (Atg8 in yeast) proteins facilitate the recruitment of cargo proteins to the phagophore by binding to LC3-interacting regions (LIRs) / Atg8-interacting motifs (AIMs) [8, 9]. In the following, we will refer to LC3 and LIR as generic names for these proteins and motifs, respectively. LC3 proteins are small proteins with a core fold similar to ubiquitin. They are modified via lipidation and anchored to the phagophore membrane [10]. They bind either directly to cargo proteins or to cargo receptor proteins, which bind to both LC3 and cargo proteins. Segments of cargo proteins binding directly to LC3s are called intrinsic receptors. Nup159, for instance, is a part of the Nuclear Pore Complex (NPC) that has been shown to interact directly with Atg8 and mediate selective autophagy of the NPC [11]. The regions in (intrinsic) cargo receptors that bind to LC3s are—typically though not exclusively—LIRs, but others, e.g., ubiquitin-interacting motif (UIM)-like sequences, which bind to LC3’s UIM-docking site (UDS), also exist[12, 13].

Canonical LIRs are short linear sequences that bind to two hydrophobic pockets in the LC3. The canonical LIR motif consists of four residues that follow the pattern Θ-X-X-Γ. Θ (=W, F or Y) binds in hydrophobic pocket 1 (HP1) and Γ (=L, I or V) in hydrophobic pocket 2 (HP2). Additionally, the LIR motif forms a parallel β-sheet with the LC3 protein [13]. Besides canonical LIRs, non-canonical LIRs have been identified that can deviate from this interaction mode in various ways [14]; e.g., they deviate from the canonical sequence motif [15], bind only one of the hydrophobic pockets [16], form a third hydrophobic interaction [17], and/or have an α-helical structure [18]. Furthermore, there are also non-continuous LIRs, in which the LC3 binding residues are distributed over a wide range of the protein sequence [19]. Canonical and non-canonical core LIR motifs are often flanked by acidic and/or phosphorylatable residues [13].

Phosphorylation modulates LC3-LIR interactions. For multiple LIRs it has been shown that their binding to LC3s can be influenced by phosphorylation of residues in close proximity to the LIR [20, 21]. Structural and binding affinity studies suggest that these phosphorylated residues interact with a range of positively charged residues in the LC3 protein and thereby stabilize the interaction. Still, in some cases phosphorylation has a negative regulatory effect [3].

A variety of experimental methods can be deployed to identify and characterize functional LIR motifs. RNAi-based screenings, Yeast-2-hybrid assays, and mass spectrometry-based proteomics are examples of tools used to identify proteins interacting with LC3s. Mutating suspected LIRs and observing the effect on binding can serve as way to identify the exact binding site(s) [7]. To show functionality in the context of selective autophagy, these motifs can be tested via *in vivo* mutation studies [11, 22]. For biophysical characterizations, e.g., by calorimetry, and structural studies, usually not the full-length protein is used. Rather, these studies are conducted with short peptide fragments containing the respective LIRs [20, 21, 23]. Computational methods complement these experimental approaches by predicting LIRs and thereby limiting the search space.

AlphaFold2 [24] has emerged as a powerful new computational tool to aid in the prediction of LIRs and to guide experimental studies. In the past, sequence-based computational methods for the prediction of LIRs were the go-to method, e.g., iLIR [25]. As possible challenges, the sequences may contain the canonical motif without being functional or be functional with non-canonical motifs. Ibrahim et al. [26] were able to show that AlphaFold2 Multimer [27] performs well in predicting LC3-LIR interactions, albeit with room for improvement. Some interactions were not detected, finding multiple LIRs in one protein requires workarounds, and the effect of post-translational modifications (PTMs) on the interactions is not considered. Alternatively, a recent data-driven approach has made use of AlphaFold2 as one part of a pipeline to identify novel selective autophagy receptors [28]. Coming from a different angle, various new approaches have been developed to make use of AlphaFold2 Multimer’s ability in the prediction of interactions between proteins and peptides / protein fragments [29, 30].

Besides LC3-LIR interactions, there are other protein-protein interactions involving short linear motifs (SLiMs) that might be targetable with similar computational approaches. Small ubiquitin-like modulators (SUMOs) are small proteins with an ubiquitin-like core fold and a short disordered N-terminal region [31]. They are linked to substrate proteins via an isopeptide bond as a PTM called SUMOylation. Proteins with SUMO interacting motifs (SIMs) can bind to SUMO’s SIM binding site [32, 33]. SUMOylation has been shown to modulate protein complex and condensate formation in multiple essential cellular processes[34]. Hence, the prediction of SIMs is of high interest as well, and has been the target of previous and recent method developments [35–37].

Here, we present a fragment-based AlphaFold2 screen to find novel LIR candidates via the prediction of LC3-LIR complexes. We scan fragments of the target protein with varying lengths for interactions with LC3 proteins. Furthermore, we also attempt to capture the effect of phosphorylations by introducing phosphomimetic mutations. We show that both fragment length and phosphomimetic mutations modulate the models created by AlphaFold2. Systematic fragment screening for a range of target proteins yields structural models for interactions that AlphaFold2 fails to predict for full-length targets. Finally, we also test the transferability of this framework by applying it to SUMO-SIM interactions.

## Results

### Target fragmentation improves AlphaFold2 predictions of LC3-LIR complexes

We found that AlphaFold2 Multimer [27] readily predicts some known LC3-LIR complex structures, but not all, consistent with earlier reports [26]. The interaction of the optineurin LIR_178_*_−_*_181_ with LC3B [20] is captured with high confidence (Fig. 1A). As evidence for advances in data and software, AlphaFold2.3 also predicted a complex between the experimentally confirmed AIM1_1078_*_−_*_1081_ of Nup159 and Atg8 [11, 23] (Fig. S1A), which was not yet captured by AlphaFold2.2. Following this trend, the recently developed AlphaFold3 consistently predicted an interaction between Joka2’s experimentally confirmed LIR [38] and ATG8CL in various “close-to-canonical” complexes (Fig. S1B), an interaction not found by AlphaFold2. By contrast, the validated interaction of calreticulin with GABARAP [39, 40] was captured neither by AlphaFold2 (Fig. 1B and S1C), as found before [26], nor by AlphaFold3 (Fig. S1D).

**Figure 1:**
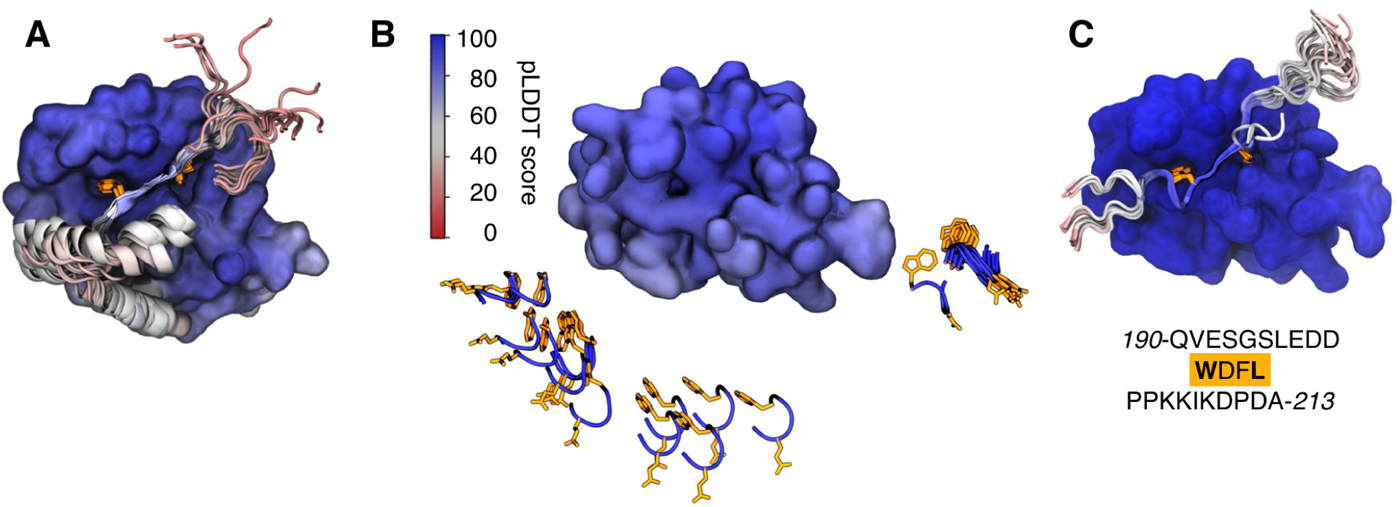
AlphaFold2 Multimer predictions with fragments can capture LC3-LIR interactions that are not found when using full-length sequences. (**A**) LC3-optineurin interaction is captured by AlphaFold2.3. Shown are 25 models of an AlphaFold2.3 prediction of the interaction between LC3B (surface representation) and optineurin (cartoon representation, with LIR residues highlighted in orange licorice), aligned on LC3B. Only the relevant part of optineurin from the respective model, and only LC3B from the top model are shown for clarity. (**B**) AlphaFold2.3 does not capture the interaction of GABARAP with full-length calreticulin. Shown are 25 models of an AlphaFold2.3 prediction of the interaction between GABARAP (surface representation) and calreticulin (cartoon representation, with LIR residues highlighted in orange licorice), aligned on GABARAP. Only the minimal LIR of calreticulin from the respective model and only GABARAP from the top model are shown for clarity. (**C**) AlphaFold2.3 captures interactions between GABARAP and a LIR-containing fragment of calreticulin. Shown are 25 models of an AlphaFold2.3 prediction of the interaction between GABARAP (surface representation) and a 24-amino-acid calreticulin fragment containing the canonical LIR (cartoon representation, with LIR residues highlighted in orange licorice), aligned on GABARAP. Only GABARAP from the top model is shown for clarity. All structures are colored by predicted local-distance difference test (pLDDT) score.

By shortening the amino-acid sequence containing a putative LIR into fragments, we expanded the set of LC3-LIR complexes captured by AlphaFold2. Experimentally, LIRs are often probed in the form of short peptide fragments, especially in structural studies [20, 21]. Therefore, we reasoned that AlphaFold2 might perform well on short peptide fragments. Indeed, it is able to predict the experimentally validated LIR in an LC3-LIR interaction for GABARAP and a 24-residue calreticulin fragment (Fig. 1C).

Varying the fragment length modulates the structural predictions of AlphaFold2, but the strength of the effect is system dependent. We systematically varied the fragment length for the LC3B-optineurin and Atg8-Nup159 systems (Fig. 2). We centered the fragments on the experimentally confirmed LIR_178_*_−_*_181_ and AIM1_1078_*_−_*_1081_ for optineurin and Nup159, respectively. The length varied from 4—just the core LIR motif—to 68 residues. The shortest segments (*<*15 residues), for which AlphaFold2 fails to generate an MSA, tend to form high-scoring canonical LC3-LIR complexes in both systems. For short to medium length fragments (approximately 15 to 40 residues), there is a weak decline in scoring for optineurin, and a strong decline for Nup159, before scores partially recover for long (*>*40 residues) fragments (Fig. 2A and B, blue lines). For Nup159 the fragment length also modulates whether a canonical or non-canonical interaction is predicted (Fig. 2C).

**Figure 2:**
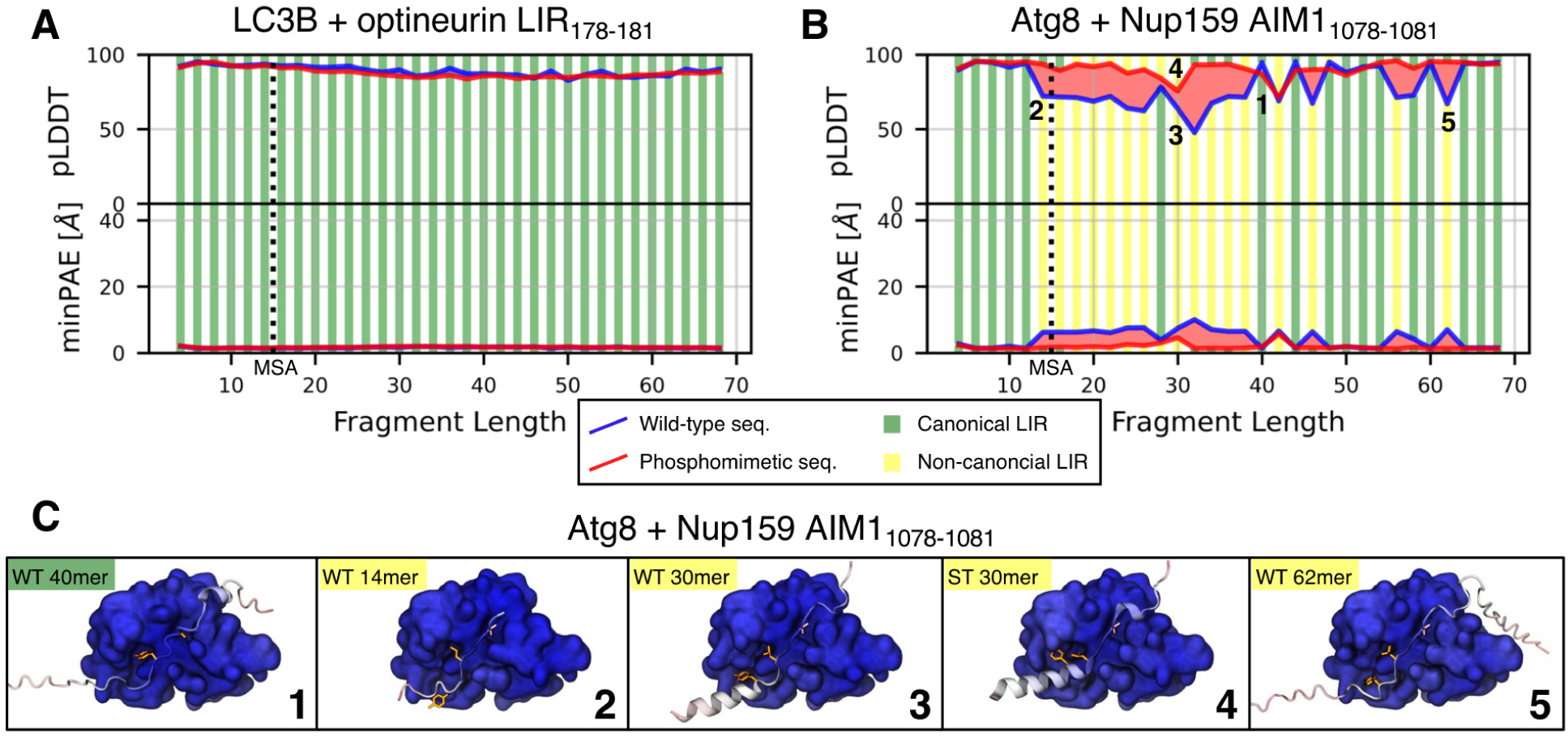
Fragment length and phosphomimetic mutations modulate the prediction of LC3-LIR interactions with AlphaFold2. Predicted local-distance difference test (pLDDT) score, minimum predicted aligned error (minPAE), and binding mode for the four core LIR/AIM residues in AlphaFold2.3 predictions of (**A**) LC3B and optineurin fragments centered on the LIR_178_*_−_*_181_ and (**B**) Atg8 and Nup159 fragments centered on the AIM1_1078_*_−_*_1081_. Fragments were varied (i) in length (x-axis) and (ii) by the introduction of phosphomimetic S/T to E mutations for experimentally confirmed phosphosites (red line) compared to wild type (WT, blue line). The vertical dotted lines indicate the minimum length required for the MSA of AlphaFold2. Stripes above the pLDDT curve / below the minPAE curve indicate the binding mode of the respective fragment with phosphomimetic mutations. Vertical bars below the pLDDT curve / above the minPAE curve indicate the binding mode of the respective WT fragment (green: canonical LIR; yellow: non-canonical LIR). (**C**) Structures for some selected fragments from **A** showing one canonical and four non-canonical LC3-LIR interactions. Nup159 fragments are shown in cartoon representation with Y1078 and L1081 highlighted in orange, and I1084 in pink licorice. Atg8 is shown in surface representation. All structures are colored by pLDDT score. Numbers in **B** correspond to structures in **C**.

### Target fragmentation and phosphorylation improve AlphaFold2 predictions of LC3-LIR complexes

Phosphorylation is a modulator of LC3-LIR interactions, with experimentally resolved structures of LC3-LIR complexes frequently containing phosphomimetic mutations or phosphorylated residues flanking the LIR [20, 21]. We therefore tested if the introduction of phosphomimetic mutations influences AlphaFold2’s prediction behavior for LC3-LIR complex structures. We introduced phosphomimetic S/T to E mutations for all S and T residues in our fragments listed in dbPTM [41], a database for PTMs. For optineurin LIR_178_*_−_*_181_ the effect on the already high scores is negligible (Fig. 2A, red lines), for Nup159 AIM1_1078_*_−_*_1081_ scores tend to improve, in some cases greatly, and seven more fragments capture the canonical interfaces (Fig. 2B, red lines). The effect is similar when using AlphaFold2.2 instead of 2.3, albeit with mostly lower model quality (Fig. S2). Overall, phosphomimetic mutations emerge as a tool to modulate the AF2 screen and identify otherwise missed LC3-LIR interactions.

### Systematic screening over multiple targets reveals known LIRs and novel LIR candidates

Based on the observed effects of fragment length and phosphomimetic mutations, we designed a systematic screen for the identification of LIR motifs (Fig. 3A). For a given target protein, we generated fragments of a defined length with 75% overlap between them. We then repeated the process with a sequence of the protein in which all S and T residues expected to be phosphorylated were mutated to E. For all resulting unique fragments, structural complexes were predicted with AlphaFold2 Multimer. Every resulting complex structure was assessed based on AlphaFold2’s quality scores and the type of interaction, similar to the fragment-length screen. To test if the predicted LIR is surface-accessible to bind to LC3, we used the AlphaFold2 predicted structure of the full-length target protein in isolation. As metrics for accessibility we calculated residue depth and secondary structure, where secondary structure was only assigned to residues with pLDDT scores above 70.

**Figure 3:**
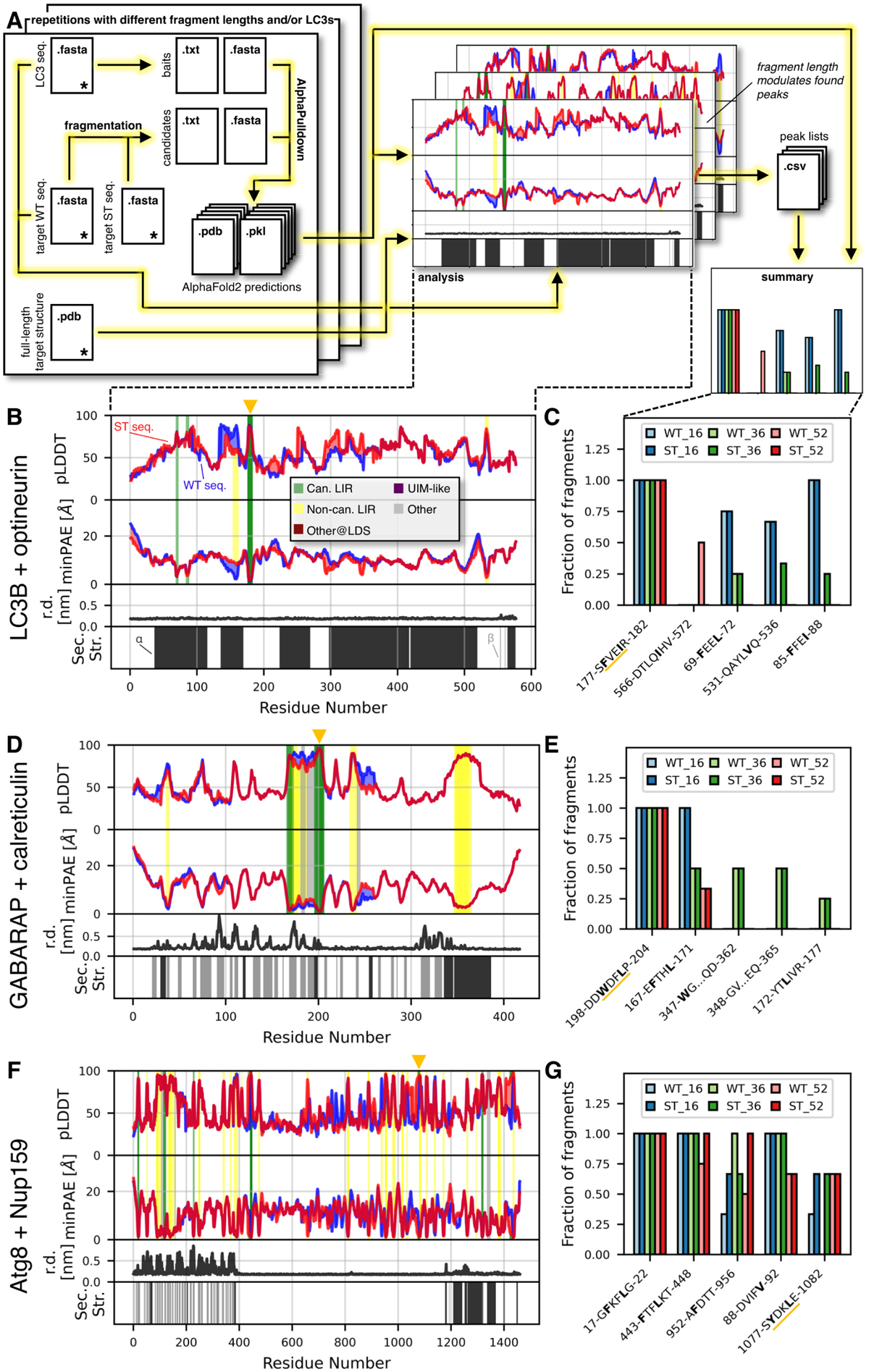
Scanning over the entire sequence of target proteins with fragment-LC3 AlphaFold2 Multimer predictions reveals known LIRs and novel candidate LIRs. (**A**) Overview over the LIR scanning pipeline. Inputs are sequences of the LC3 protein to be screened against and the wildtype (WT) target, optionally a target sequence with phosphomimetic (ST) mutations, and a structure of the target protein (all input files are marked with an asterisk). Overlapping fragments of WT and ST target sequences are then screened against the LC3 via alphapulldown [69]. The predicted complex structures are analyzed for LC3-LIR and other interactions. The input structure file provides information on the accessibility of the predicted interacting motifs in the target. Different fragment lengths provide a trade-off between sensitivity and specificity. The screen can be repeated for different fragment lengths and LC3s. The predicted binding motifs are grouped based on their similarity in a summary analysis. Relevant file formats are indicated for every step. (**B**) Systematic screen of the interaction between 36-residue fragments of optineurin with LC3B. The fragments have a 75% overlap. We performed the screen for the WT sequence, and an ST sequence, in which S/T to E mutations for experimentally confirmed phosphosites were introduced. Predicted local-distance difference test (pLDDT) score, minimum predicted aligned error (minPAE), and binding mode for the interacting residues are shown (minimum minPAE from all fragments covering each residue and its respective pLDDT score). Stripes above the pLDDT curve / below the minPAE curve indicate the binding mode of the respective fragment with phosphomimetic mutations. Stripes below the pLDDT curve / above the minPAE curve indicate the binding mode of the respective wild-type fragment. Additionally, we calculated residue depth (r.d.) and secondary structure (sec. str.; assigned only if the pLDDT score of the respective residue is *>*70) from the AlphaFold2 predicted structure for the full-length protein to add structural context to the sequence (black: α-helix; grey: β-sheet). The experimentally confirmed LIRs are indicated at the top by orange triangles. (**C**) Summary of predicted LIRs from the screen in **B** and two additional screens using the same setup, but 16residue and 52-residue fragments, respectively. Shown are the top five predicted LIRs, sorted by relative occurrence in longer fragments (52mers *>* 36mers *>* 16mers). Residues binding in hydrophobic pocket 1 and 2 are highlighted in bold. The experimentally confirmed LIR is underlined in orange. (**D and E**) Results of an analogous screen for calreticulin and GABARAP. (**F and G**) Results of an analogous screen for Nup159 and Atg8.

We performed this screen on LC3B and optineurin, GABARAP and calreticulin, and Atg8 and Nup159. For each of these systems we performed runs with short (16 residues), medium length (36 residues), and long (52 residues) fragments. We did not include very short sequences, for which no MSA can be calculated, due to a possibly increased risk for false positives. The screen found the experimentally confirmed LIRs in all three systems. Interactions between fragments and LC3s were favored for shorter fragments (Fig.S3A, B, and C), suggesting a potential trade-off between specificity and sensitivity.

We combined the information from the three screens performed for LC3B and optineurin (Fig. 3B and S3A) and grouped similar interactions (Fig. 3C). The validated LIR_178_*_−_*_181_ is consistently predicted in all fragments that contain it, for all fragment lengths, and for wild-type and phosphomimetic sequences. Furthermore, a low residue depth and no secondary structure for the motif in the full-length protein suggest that it is not buried deep in a folded domain. Similarly, the validated calreticulin LIR_200_*_−_*_203_ is found in all fragments containing it (Fig. S3B, 3D and E), and lies at the edge of a folded region, suggesting that it is partially hidden (Fig. 3D, compare to the structure shown in Fig. S1C). Nup159 AIM1_1078_*_−_*_1081_ is also reliably predicted, though not in all fragments containing it (Fig. S3C, 3F and G). For this LIR, phosphomimetic mutations substantially increase the predictive power. The fraction of fragments for which the LIR is predicted rises from 0.33 to 0.67 for 16mers and from 0.0 to 0.67 for 36mers. AIM1 is located in an unstructured segment of the protein (Fig. 3F). Contrary to optineurin and calreticulin, the experimentally confirmed LIR was not the most reliably predicted one in Nup159.

Instead, the screen suggests novel LIR candidates for some of the systems. For optineurin these alternative interacting motifs occur only in a small fraction of the fragments and lie within α-helical parts of the protein. So the evidence for them is rather weak. For calreticulin, a second LIR is reliably predicted, though it is located in a folded region and, hence, likely not available as long as calreticulin is folded. By contrast, for Nup159 the screen predicts a canonical and a non-canonical LIR, LIR_443_*_−_*_448_ (also predicted by iLIR [25]) and LIR_952_*_−_*_956_, respectively, with high reliability and in unstructured regions of the protein. Two more LIRs with strong score indication are located in the folded N-terminal domain of Nup159. As an important negative control, our list of top-scoring additional LIRs does not contain any of the previously predicted but, in experiment, non-functional AIM2, AIM3, AIM4, and AIM5 [11] in the top 5, and only one of them in the top 15 (Fig. S3D). Instead, our screen is dominated by the experimentally confirmed AIM1.

In an additional run, we tested five more systems with experimentally confirmed canonical LIR motifs. Running the screen with GABARAP against STBD1 (Fig. 4A), BNIP3 (Fig. 4B), and ULK2 (Fig. 4C) captured the experimentally confirmed [42–45]—but structurally unresolved— LIR, respectively, in all fragments. For the two remaining systems, ATG8CL with Joka2, and ATG8E with DSK2A, the results are less clear. For Joka2, the experimentally confirmed LIR [38] is predicted less consistently, with a couple of other sequences ranking clearly above it (Fig. 4D), though some of them, e.g., LIR_637_*_−_*_640_, seem to be buried in folded domains (Fig. S4A). In DSK2A, we found two of the three confirmed LIRs [46] (LIR_43_*_−_*_46_ and LIR_249_*_−_*_252_) in at least one fragment (Fig. 4E), with LIR_43_*_−_*_46_ occurring in three different binding modes (Fig. S4B), and LIR_249_*_−_*_252_ not making it into the top 15.

**Figure 4:**
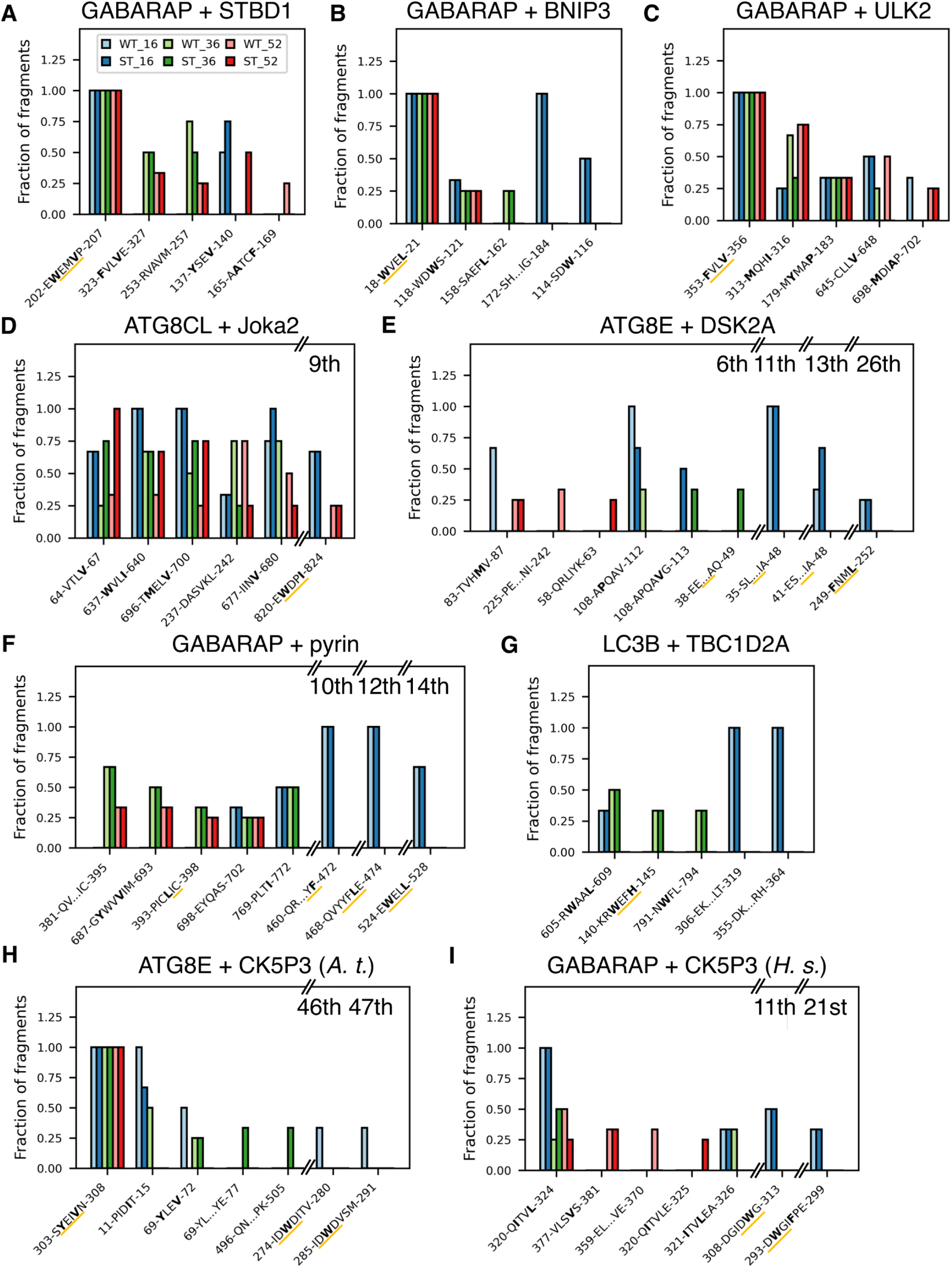
The prediction of LC3-LIR interactions produces mixed results for challenging targets. Results of fragment-prediction screens for (**A**) GABARAP and STBD1, (**B**) GABARAP and BNIP3, (**C**) GABARAP and ULK2, (**D**) ATG8CL and Joka2, (**E**) ATG8E and DSK2A, (**F**) GABARAP and pyrin, (**G**) LC3B and TBC1D2A, (**H**) ATG8E and CK5P3 (*A. t.*), and (**I**) GABARAP and CK5P3 (*H. s.*). Shown are the top five predicted LIRs and all lower-ranking predicted LIRs (rank indicated above respective bars) corresponding to experimentally confirmed ones, sorted by relative occurrence in longer fragments (52mers *>* 36mers *>* 16mers). Residues binding in hydrophobic pocket 1 and 2 are highlighted in bold. The experimentally confirmed LIRs are underlined in orange.

We also tested the screen with four very challenging targets containing multiple (non-)canonical LIRs: GABARAP and pyrin [47], LC3B and TBC1D2A [48], ATG8E and CK5P3 (*Arabidopsis thaliana*), and GABARAP and CK5P3 (*Homo sapiens*) [15]. For pyrin, our screen picked up the canonical LIR and the two non-canonical LIRs (one of them in two different binding modes; Fig. S4C), but these interactions were not reliably predicted in all fragments (Fig. 4F). For TBC1D2A, we found one of the two non-canonical LIRs, again with a rather weak signal (Fig. 4G). For CK5P3 (*A. t.*) the canonical LIR was predicted very consistently, but of the three closely grouped (Fig. S4D) non-canonical LIRs only two were predicted in at least one fragment, and none of them ranked in the top 15 (Fig. 4H). Like for Nup159, we have four negative controls—previously predicted LIRs that were experimentally verified to be non-functional—for CK5P3 (*A. t.*). Only two of these were within our top 15 candidates, also being predicted less consistently than the experimentally confirmed canonical LIR (Fig. S4E). For CK5P3 (*H. s.*), we found two of the three confirmed non-canonical LIRs, with one of them making it into the top 15 (Fig. 4I), but again with low consistency. Interestingly, the only LIR predicted consistently across fragment lengths lies in close proximity to the experimentally identified ones. Both experimental LIRs and the predicted novel candidate LIR_320_*_−_*_324_ are located in an unstructured part of the protein, while many of the other signals occur in domains expected to be folded (Fig. S4E).

### Phosphomimetic mutations mimic phosphorylation in molecular dynamics simulations

Using the Atg8-Nup159 AIM1 system as an example, we analyzed the structural effect of the phosphomimetic mutations. In molecular dynamics (MD) simulations, we compared all fragments that formed a canonical LC3-LIR interaction with Atg8 for the wild-type (Fig. 5A) and the phosphomimetic (Fig. 5B) Nup159 fragments. The core LIR residues and their C-terminal region behaved similarly. In the N-terminal region, the majority of the phosphomimetic fragments formed a short α-helix that was not observed for the wild type. This short helical stretch binds in a groove at the surface of Atg8. We hypothesized that this helix positions the phosphomimetic Nup159 residues in such a way that they can interact with Atg8.

**Figure 5:**
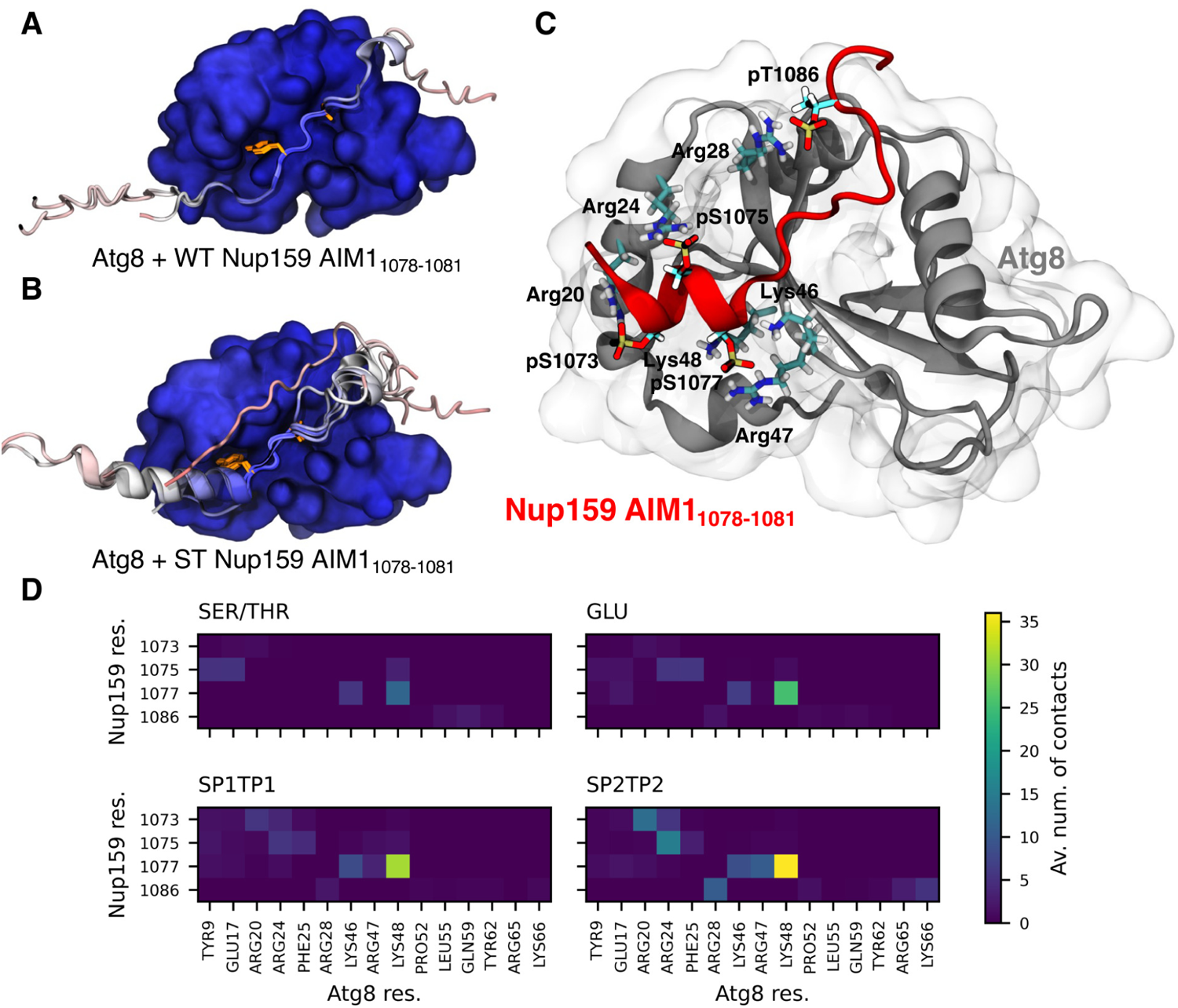
Phosphomimetic mutations alter the structure of AIM1 in Nup159, its interactions with Atg8, and behave similar to real phosphorylations. (**A**) Top models of all wild-type Nup159 fragments (cartoon representation) containing AIM1_1078_*_−_*_1081_ (Y1078 and L1081 highlighted as orange licorice) and interacting with Atg8 (surface representation) in a canonical LC3-LIR interaction. (**B**) Top models of all Nup159 fragments with phosphomimetic mutations (cartoon representation) containing AIM1_1078_*_−_*_1081_ (Y1078 and L1081 highlighted as orange licorice) and interacting with Atg8 (surface representation) in a canonical LC3-LIR interaction. In **A** and **B** only one Atg8 structure is shown for clarity, structures are aligned on Atg8, and colored by predicted local-distance difference test (pLDDT) score. (**C**) Snapshot structure of Atg8 and a Nup159 AIM1 fragment (both shown in cartoon representation) with four phosphorylations after 450 ns of MD simulation, showing some of the most prominent charge interactions. Phosphorylated residues from Nup159 and interacting residues from LC3B are highlighted in licorice. (**D**) Average number of heavy atom contacts (Av. num. of contacts; distance lower than 5 Å) per frame between potentially phosphorylated Nup159 residues and Atg8 residues. Data from triplicate (1 µs each) MD simulations for four different phosphorylation states: unphosphorylated (SER/THR), phosphomimetic mutation (GLU), and phosphorylated (SP1/TP1: -HPO_4_*^−^* and SP2/TP2: -PO_4_^2^*^−^*, respectively).

We investigated the interactions formed by the phosphorylatable residues in MD simulations of the system in different phosphorylation states. We extracted a representative structure of a 20-residue long Nup159 fragment centered on AIM1 (by truncation of a longer fragment) and bound to Atg8 for the wild-type and phosphomimetic system, respectively. For the phosphomimetic system, we also replaced the phosphomimetic residues with unprotonated (SP2 and TP2 with charge *−*2) or singly protonated (SP1 and TP1 with charge *−*1) phosphorylated residues. In all simulations, the core LIR stayed stably bound to HP1 and HP2 of Atg8, while the N- and C-terminal segments were more dynamic. This includes the short N-terminal helix, which partially unfolded in a few simulations (Fig. S5).

We found that phosphomimetic and phosphorylated Nup159 residues form similar contacts with Atg8, interacting mainly with a few positively charged residues. These residues are mainly Arg20, Arg24, Lys46, Arg47, and Lys48 for the three N-terminal phospho-residues pS1073, pS1075, and pS1077. For the C-terminal pT1086, the main interaction partner is Arg28, though we also observed some interactions with Arg65 and Lys66 (Fig. 5C and D). The general pattern of interactions is similar for phosphomimetic and phosphorylated residues, with the strength of the contacts increasing as E *<* SP1/TP1 *<* SP2/TP2. By contrast, for the unphosphorylated residues, only S1077 shows a similar pattern, while the other residues prefer different interaction partners when compared to their phosphorylated counterparts.

### Further LC3-LIR interactions from screen of LC3 proteins

We investigated the influence of the chosen LC3 protein on the predictions by AlphaFold2. Since the binding affinity between a LIR and an LC3 has been shown to vary by up to an order of magnitude for different LC3s [49], we reasoned that AlphaFold2’s predictions might reflect this. Hence, we applied our pipeline to three proteins containing LIRs with varying affinity for LC3B and GABARAP (Fig. S6).

For three examples (PLEKHM1, ULK1, and FUNDC1), we found moderate differences between LIRs predicted in interactions with LC3B and GABARAP. For the experimentally confirmed LIRs in PLEKHM1 [50] and FUNDC1 [3, 51], LIR_635_*_−_*_638_ and LIR_18_*_−_*_21_, respectively, we observed basically no difference (Fig. 6A and C). By contrast, for ULK1’s LIR_357_*_−_*_360_ [22, 45] we found a notable increase in fragments containing the interaction with GABARAP as compared to LC3B (Fig. 6B). This is consistent with the experimental data showing that the difference in affinity for ULK1’s LIR_357_*_−_*_360_ is the largest for the three screened proteins [49]. Interestingly, for other predicted LIRs with overall weaker scores, the difference between GABARAP and LC3B was larger. Whereas GABARAP tends to give stronger signals overall, the LC3B signal was found to dominate for individual LIRs. One example is the candidate non-canonical LIR_440_*_−_*_443_ for PLEKHM1. Overall, these results suggest that repeating the screen with different LC3 proteins may yield additional LIRs, and might indicate differences in affinity.

**Figure 6:**
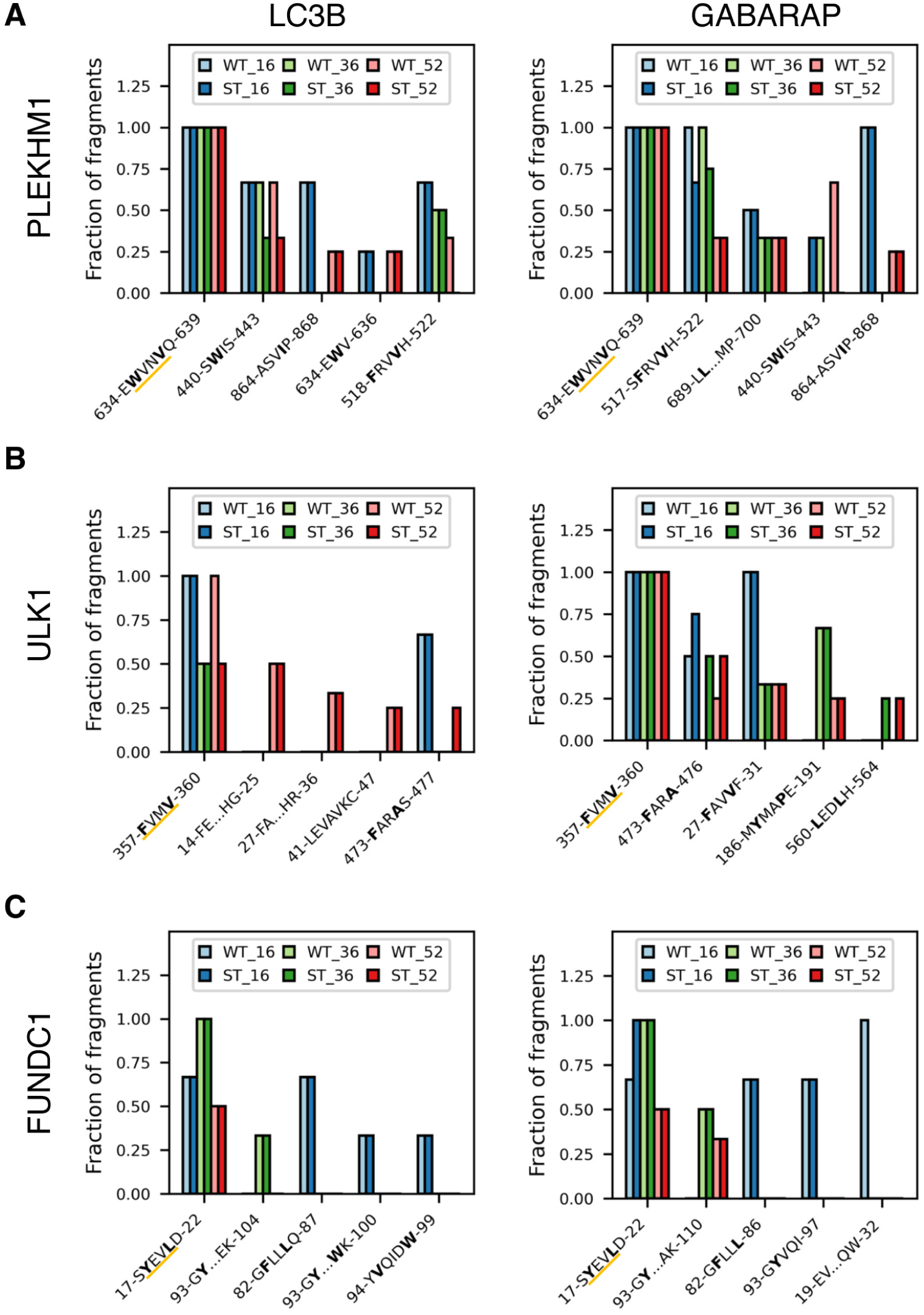
The prediction of LC3-LIR interactions depends on the LC3 protein. Results of fragment-prediction screens for (**A**) PLEKHM1, (**B**) ULK1, and (**C**) FUNDC1 combined with (i) LC3B and (ii) GABARAP. Shown are the top five predicted LIRs, sorted by relative occurrence in longer fragments (52mers *>* 36mers *>* 16mers). Residues binding in hydrophobic pocket 1 and 2 are highlighted in bold. The experimentally confirmed LIRs are underlined in orange.

### Fragment-based AlphaFold2 scan finds canonical and non-canonical candidate LIRs in NUP214

We chose NUP214 as an application example. As the human analog to Nup159 [52], NUP214 may be targeted by LC3 proteins as well. However, its length of over 2000 residues makes it a difficult target for experimental and computational screens alike.

In our NUP214 fragment screen, we found more promising candidate LIRs with GABARAP (Fig. 7B and S7A) than with LC3B (Fig. 7A and S7A). Strong LIR candidates are found in a high fraction of the fragments and are not located in well-folded parts of the protein. For the GABARAP screen, some of these candidates are sensitive to phosphomimetic mutations, including the non-canonical LIR_1560_*_−_*_1564_ and the canonical LIR_520_*_−_*_523_, LIR_1885_*_−_*_1889_, and LIR_1265_*_−_*_1268_. This last LIR is also a relatively strong and phosphorylation-dependent candidate in the LC3B screen and is predicted also by iLIR [25]. The strongest phosphorylation-dependent candidate in the LC3B screen is the non-canonical LIR_713_*_−_*_720_, which appears as LIR_709_*_−_*_718_ also in the GABARAP screen.

**Figure 7:**
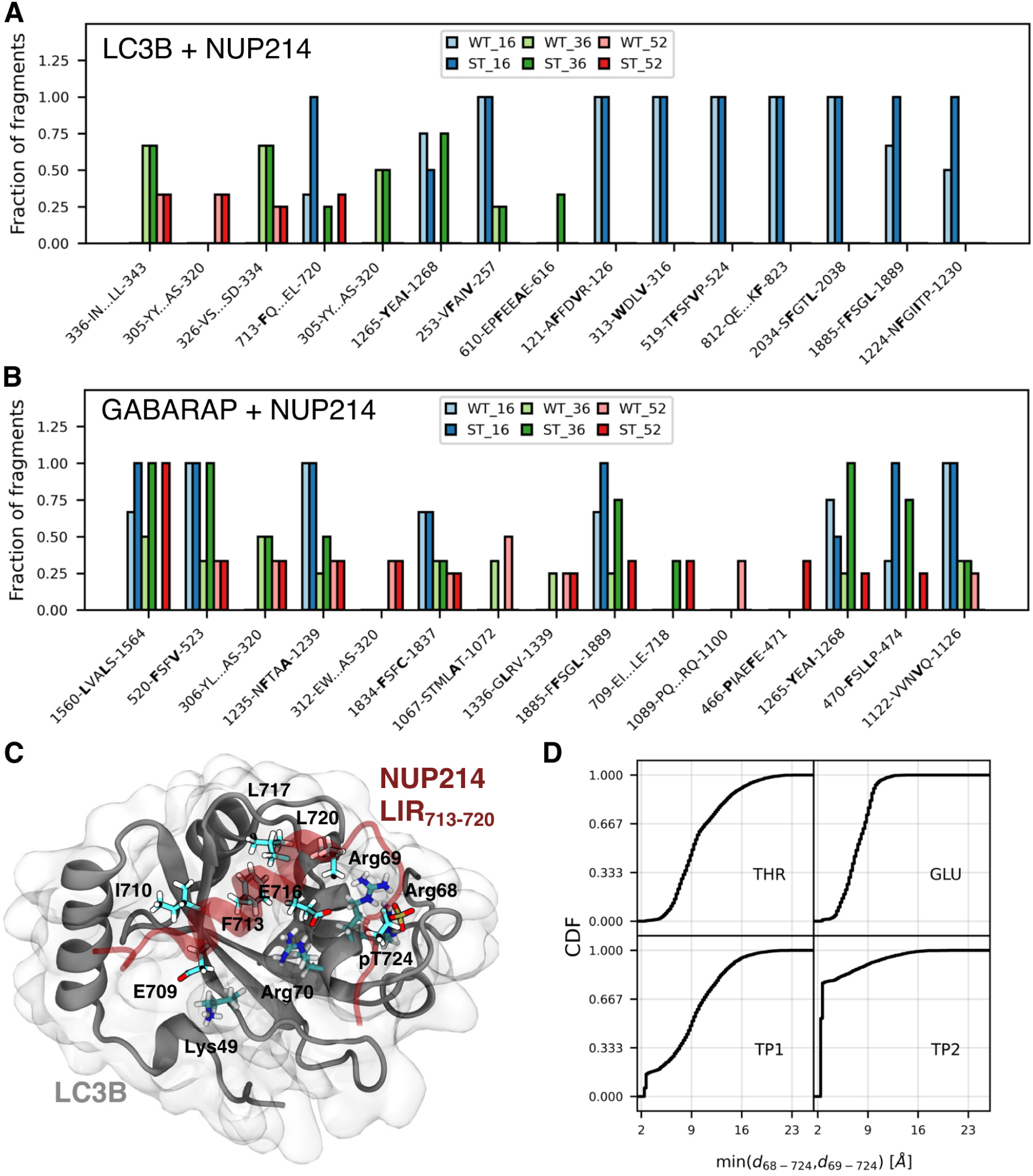
The fragment-based AlphaFold2 screen predicts multiple phosphorylation (in)dependent LIRs in NUP214. Results of fragment-prediction screens for NUP214 combined with (**A**) LC3B and (**B**) GABARAP. Shown are the top fifteen predicted LIRs, sorted by relative occurrence in longer fragments (52mers *>* 36mers *>* 16mers). Residues binding in hydrophobic pocket 1 and 2 are highlighted in bold. (**C**) Snapshot structure of LC3B and a NUP214 fragment containing the non-canonical LIR_713_*_−_*_720_ with four phosphorylations after 1 µs of MD simulation. Interface forming residues from NUP214 and some selected residues from LC3B are high-lighted in licorice. (**D**) Cumulative distribution functions (CDFs) for the minimum value of the distances between the potentially phosphorylated NUP214 residue T724 and Arg68 or Arg69 from LC3B, respectively. The two distances were computed for every timestep of the analysis and the lower one then added to the histogram. Steps below *≈*4 Å indicate the fraction of tightly coordinated structures in the MD simulations. Data from triplicate (1 µs each) MD simulations for four different phosphorylation states: unphosphorylated (THR), phosphomimetic (GLU) mutation, and phosphorylated (TP1: -HPO_4_*^−^* and TP2: -PO_4_^2^*^−^*, respectively).

To further probe these LIR candidates, we used MD simulations without and with phosphorylation. Similar to the Atg8-Nup159 AIM1 system, we found that phosphomimetic and phosphorylated NUP214 residues in LIR_1265_*_−_*_1268_ interact mainly with a few positively charged residues of LC3B and GABARAP, respectively. For the interaction with LC3B, we observed identical trends to the Nup159 system. In particular, the general pattern of interactions is similar for phosphomimetic and phosphorylated residues, with the strength of the contacts increasing from E *<* SP1 *<* SP2 (Fig. S7B). However, this trend is less clear for GABARAP, with notable differences in the interactions of SP1 and SP2 (Fig. S7C). Also, the phospho-residues interact with different residues in LC3B and GABARAP. For example, pS1257 interacts with Arg10 in LC3B, but in GABARAP Glu8 is located at this position and does not interact with pS1257. This suggests that while the general nature of these interactions is similar, they depend on the LC3 protein. The role of residues that are “charge-swapped” between LC3B and GABARAP has been discussed previously [53] and phosphorylation might add an additional layer contributing to LC3 specificity.

LIR_713_*_−_*_720_ forms an α-helix that dynamically interacts with LC3B via a hydrophobic interface and pT724. In the MD simulations, the predicted helical LIR segment stayed bound to LC3B for 1 µs independent of T724’s phosphorylation state (Fig. S7D). It forms a hydrophobic interface with LC3B with the residues I710, F713 (binds HP2), L717, and L720. E709 and E716 further stabilize this interface by interacting with Lys49 and Arg70 (Fig. 7C). However, the hydrophobic interface is quite dynamic, with the helix displaying varying conformations and partial unbinding (Fig. S7D). Replacing T724 by a phosphomimetic or phosphorylated residue strengthens the interaction with Arg68 and Arg69. In the fully deprotonated state (TP2 with charge *−*2), pT724 is firmly anchored to these residues (Fig. 7D). The dynamic and not completely stable binding of the helix, and the firm anchoring of pT724 suggest a strong phosphorylation dependence of this interaction, consistent with the strong dependence of the AlphaFold2 prediction on phosphomimetic mutations for this candidate LIR.

### Fragment-based AlphaFold2 screen of SLiM-mediated interactions

We tested the transferability of our method to other interactions mediated by SLiMs, focusing on SUMO-SIM interactions as an example. Similarly to canonical LIRs, SIMs are SLiMs and their binding to SUMO proteins can be modulated by phosphorylation [33, 54]. As poof of principle, we studied the complex between SUMO2 and the SIM_40_*_−_*_43_ of RAP80 [55](Fig. 8A).

**Figure 8:**
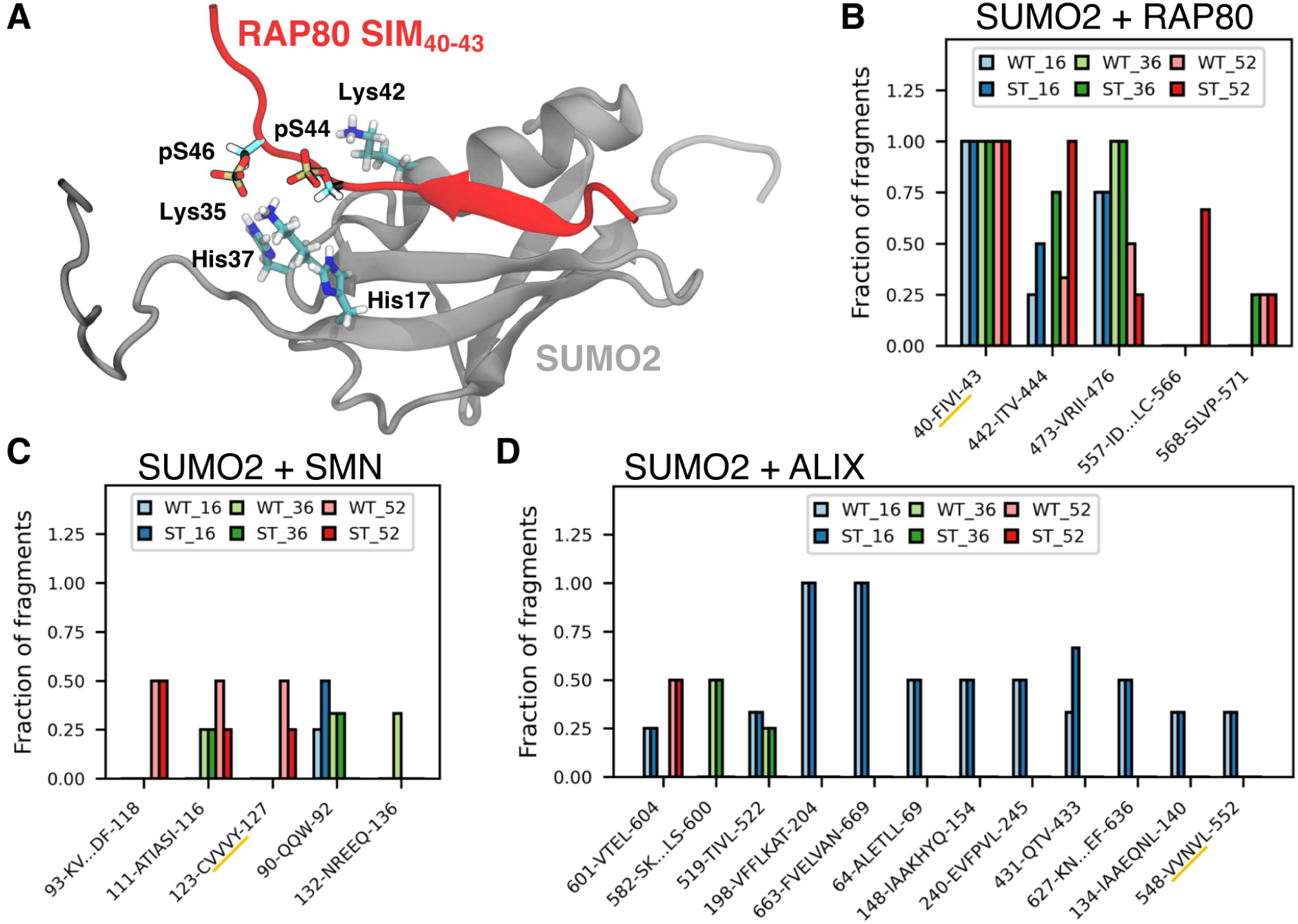
Fragment-based predictions may be suitable for other protein-protein interactions involving short linear motifs, including SUMO-SIM interactions. (**A**) Previously determined NMR structure of the RAP80-SIM bound to SUMO2 (PDB ID: 2N9E [93]). Phosphorylated residues from RAP80 and interacting residues from SUMO2 are highlighted as licorice. Results of fragment-prediction screens for (**B**) RAP80, (**C**) SMN, and (**D**) ALIX combined with SUMO2. Shown are the top five (twelve for ALIX) predicted LIRs, sorted by relative occurrence in longer fragments (52mers *>* 36mers *>* 16mers). The experimentally identified SIMs are underlined in orange.

Screening RAP80 fragments with and without phosphomimetic mutations returns the experimentally confirmed SIM in all fragments (Fig. 8B). Additionally, there are two more SIMs predicted with high consistency: SIM_442_*_−_*_444_ and SIM_473_*_−_*_476_. Interestingly, the experimentally confirmed SIM_40_*_−_*_43_ and SIM_473_*_−_*_476_ both follow the canonical Φ-Φ-X-Φ / Φ-X-Φ-Φ SIM motif, where Φ is I, V or L, and X any amino acid [32], and their prediction is not—or only very weakly—influenced by phosphomimetic mutations. By contrast, the prediction of the candidate non-canonical SIM_473_*_−_*_476_ strongly depends on these mutations. The experimentally confirmed SIM and the two strong candidates are all located in relatively unstructured regions of the protein that are accessible to interactions (Fig. S8A).

For two other proteins, finding the experimentally identified SIMs proved more challenging. We applied our pipeline to SMN and ALIX, two proteins for which recently SIMs have been identified via mutation studies [56, 57]. However, no complex structure between the respective SIM and SUMO2 was determined, making them interesting targets for our computational screen. For both systems, the fraction of fragments containing predicted SIMs is noticeably lower than, e.g., for RAP80. Nonetheless, the experimentally identified SIM_124_*_−_*_127_ is the third strongest signal we found for SMN (Fig. 8C). For ALIX, the experimentally identified SIM_548_*_−_*_552_ is only at the 12th position (Fig. 8D). Strikingly though, for SMN and ALIX the top scoring SIM candidates are all in close proximity to the ones experimentally confirmed by mutations. It is also notable that for both SMN and ALIX the experimentally identified and the strongest predicted SIM candidates lie within or at the edge of folded domains. Still, the low residue depth for these motifs suggests some degree of accessibility (Fig. S8B and C).

## Discussion

Computational prediction methods are powerful tools to design and guide experimental studies. The identification of functional—canonical and non-canonical—LIRs is a critical step in the identification of cargo and cargo receptor proteins in selective autophagy. Computational methods help to design and guide experimental studies aimed at identifying new LIRs. Here, we investigated the feasibility of a fragment-based approach that makes use of the predictive power of AlphaFold2 to find novel LIR candidates via the prediction of LC3-LIR complexes (results summarized in Table S1).

We reasoned that a fragment-based screen could be an alternative or complimentary approach to a full-length prediction. Obvious advantages of such an approach are its ability to find multiple LIRs for a sequence (e.g., for Nup159, Fig. 3G), and hidden LIRs, which would be (partially) occluded in the complete structure and become accessible only upon (local) unfolding (e.g., for calreticulin, Fig. 1B and C). Possible disadvantages are a likely higher number of false positives, in particular of buried LIRs that are unavailable for interaction in the protein in functional conformations. This problem can be addressed by including structural context like residue depth and secondary structure, since reasonable models of the full-length structures (of only the protein, not the complex) are readily available thanks to AlphaFold2 and 3. Still, fragmentation decreases the chance of finding discontinuous LIRs, and due to the multiple predictions needed for a single protein, a fragment-based screen is also computationally more expensive. Despite these drawbacks, the example of calreticulin highlights quite nicely that the additional sensitivity of this approach is essential in some cases. Also, once very long proteins are involved, full-length complex structure predictions become difficult or even unfeasible. Combined, this results in a range of cases in which a fragment-based approach is preferable to a full-length prediction.

The length of the fragments plays an important role in the search for LIRs and other SLiMs. Altering the length of fragments may be inconsequential in some cases, e.g., for the LC3Boptineurin system. In other cases, e.g., the Atg8-Nup159 system, fragment length has a strong effect on the quality of the prediction. Furthermore, in our sequence scans we found that shorter fragments tend to produce notably more protein complex structures than longer ones. This indicates a possible trade-off between false positives and false negatives depending on fragment length. However, this relation does not hold strictly true in all cases, and some interactions are only picked up in longer fragments, most notably for the SUMO2-SMN system. Repeating the screen with various fragment lengths offers a good compromise between sensitivity and specificity, albeit at increased computational cost.

The phosphomimetic mutations we introduced seem to have an overall neutral or positive effect on the prediction of LC3-LIR interfaces. Phosphomimetic mutations do not meaningfully influence the prediction of experimentally confirmed LIRs in many of the tested cases, e.g., for optineurin, calreticulin, PLEKHM1, ULK1, and FUNDC1. However, they played a critical role in finding Nup159’s AIM1 (Fig. 3G), and many of the novel LIR or SIM candidates identified in, e.g., ULK1, NUP214, and RAP80. Interestingly, binding of optineurin’s LIR to LC3B has been shown to be strongly phosphorylation dependent in experiments [20]. This indicates that while the effect of phosphomimetic mutations on the predicted interaction might serve as an indication for true phosphorylation dependence, it may not show up in the screen because the prediction is already at a very high confidence for the wild type.

It should be noted that in some of the very well performing cases this might be related to the systems having structures available, and hence being part of AlphaFold2’s training set. While the screen does perform extremely well for these cases, it does also yield good results for LIRs for which the structure is not yet solved, e.g., for ULK2, STBD1, and AIM1_1078_*_−_*_1081_ in Nup159. Furthermore, the confidence of the prediction in some of the novel candidates rivals the one for the structurally confirmed LIRs. The case of calreticulin highlights that even with a structure for the interaction being available, it is no guarantee that AlphaFold2 will predict it given the fulllength protein sequences. Hence, we argue that the screen is capable of not only reproducing existing structures, but also of generating high-confidence predictions of novel LIR candidates. Notably, in many cases where the screen produced only weak results for experimentally confirmed interactions, it found stronger signals close by, e.g., for GABARAP-CK5P3 (*H. s.*), SUMO2-SMN, and SUMO2-ALIX. On one hand, these signals may just be false positives and examples for the limitation of this method. On the other hand, there are two additional interpretations: First, the experimental results for these systems might be incomplete, as they are based on mutation studies without structural evidence. Hence, it is possible that these mutations do not lie in the actual SLiM but indirectly affect the interaction with the proximal true interface. Second, the identified motifs may offer interactions in addition to the experimentally identified ones that lead to an increase in avidity and may also increase specificity [58]. This is especially intriguing for proteins binding SUMO2, since it forms poly-SUMO2 chains in multiple cellular contexts. These SUMO2 chains provide the complementary multivalent binding interfaces for proteins with multiple SIMs [59]. Also, it has been recently proposed that in the context of poly-SUMO multivalency, auxiliary interactions not involving SIMs indeed play a role [60]. Similar to SUMO chains, one can think about multiple LC3s attached to a phagophore as multivalent binding spots for proteins with multiple LIRs, as has been discussed previously [61].

The screen is also capable of picking up differences in the interaction between a LIR and different LC3s if they are large enough. PLEKHM1’s and ULK1’s experimentally identified LIRs have been found to interact more strongly with GABARAP [49]. Our screen picked this up for ULK1, where the difference in binding affinity is the largest. For FUNDC1’s LIR_18_*_−_*_21_, a slight preference for LC3B is expected, but not picked up by our screen. Similarly, the slight preference of PLEKHM1’s LIR_635_*_−_*_638_ for GABARAP is not picked up. For NUP214, we found more strong signals when screening against GABARAP than when screening against LC3B, though individual signals, e.g., for non-canonical LIR_713_*_−_*_720_ can be stronger for LC3B. Overall, screening against GABARAP tends to generate more strong-signal candidate LIRs.

We used MD simulations to investigate the interactions of phosphorylations in LIR binding. For the two canonical LIR motifs we simulated, AIM1_1078_*_−_*_1081_ in Nup159 and candidate LIR_1265_*_−_*_1268_ in NUP214, basic residues interact with the phospho-sites. Interestingly, the interaction pattern in the simulations of GABARAP with NUP214’s LIR_1265_*_−_*_1268_ deviates slightly for the singly protonated SP1 system, highlighting some strong interactions with Leu55, Thr56, and Gln59. Visual inspection reveals that these are formed by H-bonds between the phosphate group of residue pS1271 and the backbone and sidechains of Thr56 and Gln59. Notably, phospho-residue interactions with most of the mentioned positively charged residues (or their analogs in other LC3 structures) have been described previously for other LC3-LIR interactions [20, 21, 62]. The one non-canonical LIR we simulated, candidate LIR_713_*_−_*_720_ of NUP214, contains only one phosphorylation, which interacts strongly with Arg68 and Arg69 of LC3B. Additionally, residues Lys49 and Arg70 form interactions with acidic residues flanking the hydrophobic LC3-LIR interface. For all LC3-LIR interactions we simulated, the additionally formed contacts of phosphomimetic and phosphorylated residues are consistent with the increased fraction of fragments that contain the predicted LIR, once phosphomimetic mutations for selected serine and threonine are introduced in the AlphaFold2 screen.

We did not include phosphomimetic mutations of tyrosines for two reasons. Like serine and threonine, tyrosines are also commonly phosphorylated, and have also been shown to modulate LC3-LIR interactions [3]. However, since tyrosines can serve as the Θ residue for LIRs, phosphorylated tyrosines are more likely to have a negative effect on LC3-LIR interactions, making them less useful when the goal is to identify novel LIRs. Additionally, while phosphomimetic mutations of serine and threonine to glutamate are structurally similar, for tyrosines the changes would be quite substantial. Still, if one’s priority is to assess the effect of phosphorylation on the interaction, tyrosine phosphorylation should be considered, e.g., by including only modifications for residues targeted by one specific kinase. Here, very recent developments allowing the explicit and direct consideration of PTMs in protein structure predictions, as in the recently published AlphaFold3 [63], should prove valuable.

### Recommendations for practitioners

In LIR-screens with limited compute budget, we recommend to extend the screen to the fulllength target protein, to 36mer fragments of the relevant regions of the target protein, and then to repeat the process with phosphosites (S, T) changed to phosphomimetic E. As outlined in Fig. 3A and supported by the results in Figs. 3B-G and S3A-C, this fragment length offers a good compromise between specificity and sensitivity at limited computational cost. By testing their accessibility in experimental or AlphaFold structures of the intact target protein, the list of candidate LIRs can be further trimmed. It is also possible to assess the full structure in advance and exclude well-folded segments from the screen to reduce computational cost. However, this may leave partially buried LIRs, as in calreticulin, undiscovered. A detailed examination of the predicted structures for strong candidates and of the AlphaFold indicators of confidence in the predicted interactions then gives further insights into the expected strength of the LIR predictions obtained from the computational screen and a basis to select peptides and mutation sites for experimental validation.

## Concluding remarks

With AlphaFold3 being available as an online server now [63], we expect further future improvement of our pipeline. Among its novel features, the possibility to include real PTMs in the structural predictions makes not only phosphotyrosines more accessible, but also allows the inclusion of, e.g., acetylation. Thus, one can include a greater variety of PTMs that have been shown to modulate LC3-LIR [61] and SUMO-SIM [64] interactions. Also, we observed a general improvement of AlphaFold2’s predictions between versions 2.2 and 2.3, and also—though still anecdotal—between versions 2.3 and 3. A continuation of this trend would be welcome both for fragment-based but also for full-length-based LIR predictions. Still, with the source code unavailable, and the current resource and terms-of-use based limitations, AlphaFold3 is at the moment not feasible for high-throughput peptide screens, though that may change in the future. Besides AlphaFold3, there are also other approaches on the way to better account for phosphorylation in AlphaFold2-based predictions [65]. Additionally, our approach can easily be adapted for other structure prediction methods, such as OpenFold [66] and RoseTTAFold2 [67], though we did not test how this will affect results. Lastly, as an increased number of noncanonical LC3-LIR structures will become available in the future, one can expect the quality of predictions of these complex structures to improve further.

## Methods

### Fragment sampling pipeline

#### Generation of fragment sequences

We employed two different schemes for the generation of fragment sequences, one to investigate the effect of fragment length and another to screen over an entire protein sequence. For the effect of fragment length, we chose a 4-residue core LIR as the minimum sequence. We increased the length in 2-residue steps (+1 at each terminus) up to a final length of 68 residues (32 N-terminal residues, 4 core LIR residues, 32 C-terminal residues). For the screening over protein sequences, we generated fragments of a defined length (52mers, 36mers, and 16mers) that jointly cover the entire protein sequence with 75% overlap. E.g., for 16mers, the first fragment contained residues 1-16, the second 5-20, the third 9-24, etc. For the last fragment, the starting residue was shifted, if necessary, to achieve the required length. We took sequences for all proteins from UniProt [68] (Table 1), and experimental phosphorylation sites from dbPTM [41] (with the exception of pyrin and TBC1D2A, for which we used the PTM data from UniProt). For sequences, for which no large-scale data on experimental phosphosites were available, we assumed all S and T residues to be potential phosphosites.

**Table 1:** Protein sequences used in this study.

| Protein name | UniProt ID |
| --- | --- |
| Atg8 | P38182 |
| ATG8CL | M1C146 |
| ATG8E | Q8S926 |
| LC3B | Q9GZQ8 |
| GABARAP | O95166 |
| optineurin | Q96CV9 |
| calreticulin | P27797 |
| Nup159 | P40477 |
| Joka2 | M1BJF6 |
| BNIP3 | Q12983 |
| STBD1 | O95210 |
| ULK2 | Q8IYT8 |
| pyrin | O15553 |
| TBC1D2A | Q9BYX2 |
| DSK2A | Q9SII9 |
| CK5P3 (*Homo sapiens*) | Q96JB5 |
| CK5P3 (*Arabidopsis thaliana*) | Q9FG23 |
| PLEKHM1 | Q9Y4G2 |
| ULK1 | O75385 |
| FUNDC1 | Q8IVP5 |
| NUP214 | P35658 |
| SUMO2 | P61956 |
| RAP80 | Q96RL1 |
| SMN | Q16637 |
| ALIX | Q8WUM4 |

#### Fragment and LC3 complex structure prediction

We used alphapulldown (versions 0.22.3 and 0.30.7 with AlphaFold2.2 and 2.3, respectively) [69] for the high-throughput AlphaFold2 Multimer [27, 70] predictions. The only default setting we changed was the number of cycles (ncycles), which we increased to ten. We set the “max_template_date” parameter to “2050-01-01”. We treated every fragment as an individual protein, meaning we calculated a multiple sequence alignment (MSA) for every single one. If not stated otherwise, AlphaFold2 refers to version 2.3.

#### Evaluation of generated structures

We evaluated the generated structures in two steps. The first step differed between the scans of fragment length and over entire sequences. For the length scans, we calculated the AlphaFold2 confidence scores for the four core LIR residues the respective scan was centered on. When scanning over the entire protein sequence, we looked for any interface that AlphaFold2 predicted with confidence. In the second step, we classified the confidently predicted structures based on the type of interaction between LC3 (or SUMO) and the respective fragment.

We used AlphaFold2’s “predicted local-distance difference test” (pLDDT) score and predicted aligned error (PAE) to quantify AlphaFold2’s confidence in its prediction. For each individual residue, we used the pLDDT score as given by AlphaFold2 and calculated a minimum PAE (minPAE) as the average of the five lowest PAE values for the residue. We looked for stretches in fragments, which fulfilled three criteria: (i) a minimum length of 4 residues, (ii) a minimum average pLDDT of 75 over all stretch residues, and (iii) a maximum average minPAE of 4.0 Å.

We classified LC3-stretch interactions based on their structure either as “canonical LIR” (can. LIR), “non-canonical LIR” (non-can. LIR), “other interaction proximal to the LIR-docking site” (other@LDS), interacting via an “ubiquitin-interacting motif-like” (UIM-like) sequence, or “other”. For the classification of canonical LIRs, we used four criteria: (i) Binding of a residue to HP1, (ii) binding of a residue to HP2, (iii) two or more backbone H-bonds with the LC3 β-sheet between HP1 and HP2, and (iv) the stretch having the canonical LIR sequence Θ-X-X-Γ—where Θ is W, F or Y, and Γ is L, I or V—with Θ having to be the residue bound in HP1 and Γ the one bound in HP2. Only if all four of these criteria were fulfilled, we classified a stretch as canonical LIR. If the stretch satisfied any one to three (but not all four) of these criteria, we classified it as non-canonical LIR.

We determined binding to HP1 and HP2 by evaluating all residues from N- to C-terminus of the stretch until we found one that fulfilled certain distance constraints. We considered a residue bound to HP1 if the distance between any of its heavy atoms and two defined LC3 C_α_ atoms was below a given threshold. The first distance to the C_α_ atom of residue 52 (LC3B) / 51 (ATG8E) / 50 (ATG8CL) / 49 (GABARAP and Atg8) had a cutoff of 5.25 Å, the second distance to the C_α_ atom of residue 108 (LC3B) / 106 (ATG8E) / 105 (ATG8CL) / 104 (GABARAP and Atg8) had a cutoff of 9.75 Å. For HP2, we used a similar approach with two different respective distances. The first distance to the C_α_ atom of residue 54 (LC3B) / 53 (ATG8E) / 52 (ATG8CL) / 51 (GABARAP and Atg8) had a cutoff of 4.5 Å, the second distance to the C_α_ atom of residue 67 (LC3B) / 66 (ATG8E) / 65 (ATG8CL) / 64 (GABARAP and Atg8) had a cutoff of 9.50 Å.

We used the dssp [71] definition with the default cutoff of *−*0.5 kcal/mol to determine H-bonds. Since the structures generated with alphapulldown lacked hydrogen atoms, we added backbone hydrogens in post-processing via PyMol’s h_add [72] method. We calculated the number of H-bonds between all residues of the confidently predicted stretch and the LC3 β-sheet between HP1 and HP2: residues 50-55 for LC3B, residues 49-54 for ATG8E, residues 48-53 for ATG8CL, and residues 47-52 for Atg8 and GABARAP.

All stretches without a LIR were classified as “other interaction proximal to the LIR docking site” if they formed at least five C_α_ atom contacts (distance below 5 Å) with the LC3 β-sheet between HP1 and HP2. Stretches with at least five C_α_ atom contacts with the LC3 UIM-like docking site (UDS) (residues 79-82 for LC3B, residues 78-81 for ATG8E, residues 77-80 for ATG8CL, and residues 76-79 for Atg8 and GABARAP) received a “UIM-like” classification. All remaining stretches were classified as “other”.

For SUMO2-SIM interactions, we made no distinction between canonical and non-canonical SIMs. We counted backbone H-bonds between the β-sheet of SUMO2’s SIM binding groove (residues 29-34) and the interacting confidently predicted stretch. For two or more H-bonds, we classified the interaction as a SUMO-SIM interaction. All other interactions we summarized as “other”.

#### Estimation of residue occlusion

We used residue depth with respect to the protein surface and local secondary structure as quantifiers of residue occlusion in structures of the full-length target sequence. To get full-length structures we used AlphaFold2 (see section “AlphaFold2 predictions of full-length sequences”). We calculated residue depth via msms [73] and secondary structure via dssp [71]. We used the simplified definition for structural elements and required a residue pLDDT of at least 70 to assign secondary structure.

#### Summary of predicted interaction motifs

We grouped stretches predicted in different fragments of varying length with and without phosphomimetic mutations together, if they predicted the same interaction type (e.g., canonical LIR or non-canonical LIR) and had an overlap of at least 3 residues. We ordered the summarized predictions by the relative occurrence of the interaction, meaning the number of fragments that contain the sequence stretch and form the interaction divided by the number of all fragments containing it. We calculated this property for each fragment length and mutation state individually. We ordered the predicted interacting stretches by the sum of this relative occurrence score for a given fragment length, giving longer fragments priority over shorter ones (52mers *>* 36mers *>* 16mers).

### AlphaFold2 predictions of full-length sequences

We used local installations of AlphaFold2.2 and AlphaFold2.3 Multimer, as well as the AlphaFold3 online server [63], to predict complex structures of full-length proteins from their amino acid sequence. For single proteins we used already predicted structures provided by the AlphaFold Protein Structure Database [74] if available, and AlphaFold2.3 otherwise. If not stated otherwise, AlphaFold2 refers to version 2.3.

### Molecular dynamics simulations

We used Gromacs (version 2023.2 for system preparation and equilibration, version 2023.3 for production runs) [75] and the charmm36m (July 2021 version) force field with an increased Lennard-Jones ɛ parameter for the water hydrogens for all molecular dynamics (MD) simulations [76]. We ran simulations of Atg8 in complex with Nup159 residues 1070-1089 (AIM1_1078_*_−_*_1081_), LC3B in complex with NUP214 residues 1257-1272 (LIR_1265_*_−_*_1268_), LC3B in complex with NUP214 LIR_1265_*_−_*_1268_, and GABARAP in complex with residues 703-728 (LIR_713_*_−_*_720_). All initial complex structures came from AlphaFold2 predictions in the context of the screen, with some being truncated. For AIM1_1078_*_−_*_1081_ and LIR_1265_*_−_*_1268_ systems we used predictions with wild-type sequences for wild-type simulations and predictions with phosphomimetic sequences for simulations of phosphomimetic/phosphorylated structures. For LIR_713_*_−_*_720_ systems we used a prediction based on a phosphomimetic sequence also for wild-type simulations by reversing the phosphomimetic mutations via a custom Python script. We used charged termini for LC3 proteins and N-terminal acetylation and C-terminal aminomethylation capping groups for LIR-containing fragments. Phosphorylations were introduced into the structures with phosphomimetic mutations (replacing them) via a custom Python script. We solvated the systems in TIP3P water and added 150 mM NaCl plus neutralizing ions.

We minimized all systems using the steepest descent algorithm with maximum force for convergence of 1000 kJ mol*^−^*^1^ nm*^−^*^1^. Equilibration was done with one NVT and two NPT runs, all using a timestep of 1 fs, running for 1 ns, 1 ns, and 5 ns respectively. The NVT runs used a Berendsen thermostat [77] to maintain a temperature of 300 K with a characteristic time τ_T_ of 0.1 ps. The first NPT run used a Berendsen thermostat to maintain a temperature of 300 K with τ_T_ = 0.1 ps, and an isotropic Berendsen barostat [77] with a target pressure of 1 bar, a characteristic time τ*_p_* of 5.0 ps, and a compressibility of 4.5 *·* 10*^−^*^5^ bar*^−^*^1^. The second NPT run used a v-rescale thermostat [78] to maintain a temperature of 300 K with τ_T_ = 1.0 ps, and an isotropic Parrinello-Rahman barostat [79] with a target pressure of 1 bar, a τ*_p_* of 5.0 ps, and a compressibility of 4.5 *·* 10*^−^*^5^ bar*^−^*^1^. The minimization and the first two equilibration steps used position restraints on all protein heavy atoms.

Production runs were performed for 1 µs with a timestep of 2 fs. For temperature coupling we used a v-rescale thermostat with a target temperature of 300 K, and τ_T_ = 1 ps. For pressure coupling we used an isotropic Parrinello-Rahman barostat with a target pressure of 1 bar, τ*_p_* = 5.0 ps, and a compressibility of 4.5 *·* 10*^−^*^5^ bar*^−^*^1^.

All simulations used a leap-frog integrator, a Verlet cutoff-scheme [80], a pair-distance cutoff (1.2 nm) with a force-switch modifier (1.0 nm) for Van-der-Waals interactions, a cutoff of 1.2 nm for Coulomb interactions, and Particle Mesh Ewald for long-range electrostatics [81]. Bonds for hydrogens were turned into constraints and treated with the LINCS algorithm [82].

We determined the average number of contacts for every residue pair by calculating the distances between their heavy atoms. If any pair distance was below 5 Å, the two residues were counted as being in contact. We averaged over all frames (in 1 ns timesteps) of triplicate simulations. Plots show the 15 LC3 residues with the highest average number of contacts summed over the shown target residues. We calculated the secondary structure in simulation trajectories via dssp [71] in timesteps of 1 ns.

### Implementation and general analysis

We visualized all structures with VMD [83], which was also used for structural alignment and general analysis. Furthermore, we employed Python3 [84] with Anaconda3 [85], iPython [86], Numpy [87], Matplotlib [88], Seaborn [89], MDAnalysis [90], MDTraj [91], BioPython [92], and PyMOL [72] to implement the analysis described in the previous sections.

## Abbreviations

AIM: Atg8-interacting motif
Av. num. of contacts: average number of heavy atom contacts
A. t.: Arabidopsis thaliana
can. LIR: canonical LIR
CDF: cumulative distribution function
HP1/2: hydrophobic pocket 1/2
H. s.: Homo sapiens
LDS: LIR-docking site
LIR: LC3-interacting region
minPAE: minimum PAE
MSA: multiple sequence alignment
non-can. LIR: non-canonical LIR
NPC: Nuclear Pore Complex
other@LDS: other interaction proximal to the LIR docking site
PAE: predicted aligned error
pLDDT: predicted local-distance difference test
PTM: post-translational modification
seq.: sequence
SIM: SUMO-interacting motif
ST: phosphomimetic
SUMO: small ubiquitin-like modifier
SLiM: short linear motif
UDS: UIM-docking site
UIM: ubiquitin-interacting motif
WT: wild type

## Acknowledgements

We thank Jenny Sachweh, Marcel Heinz, Jürgen Köfinger, Martin Beck, Roberto Covino, and Volker Dötsch for insightful discussions and support. We thank the Max Planck Society, the Clusterproject ENABLE funded by the Hessian Ministry for Science and the Arts, and the Collaborative Research Center 1177 “Molecular and functional characterization of selective autophagy” funded by the Deutsche Forschungsgemeinschaft (SFB1177, DFG Project-ID 259130777), for financial support, and the Max Planck Computing and Data Facility for computational resources.

## Data and code availability

All data of this study (lists of all predicted LIRs, AlphaFold structures, MD simulation trajectories) are available from the authors upon reasonable request. The code used to setup the alphapulldown runs and analyze the resulting AlphaFold predictions is publicly available under https://github.com/bio-phys/af2_lir_screen.

## Supplementary information

**Figure S1:**
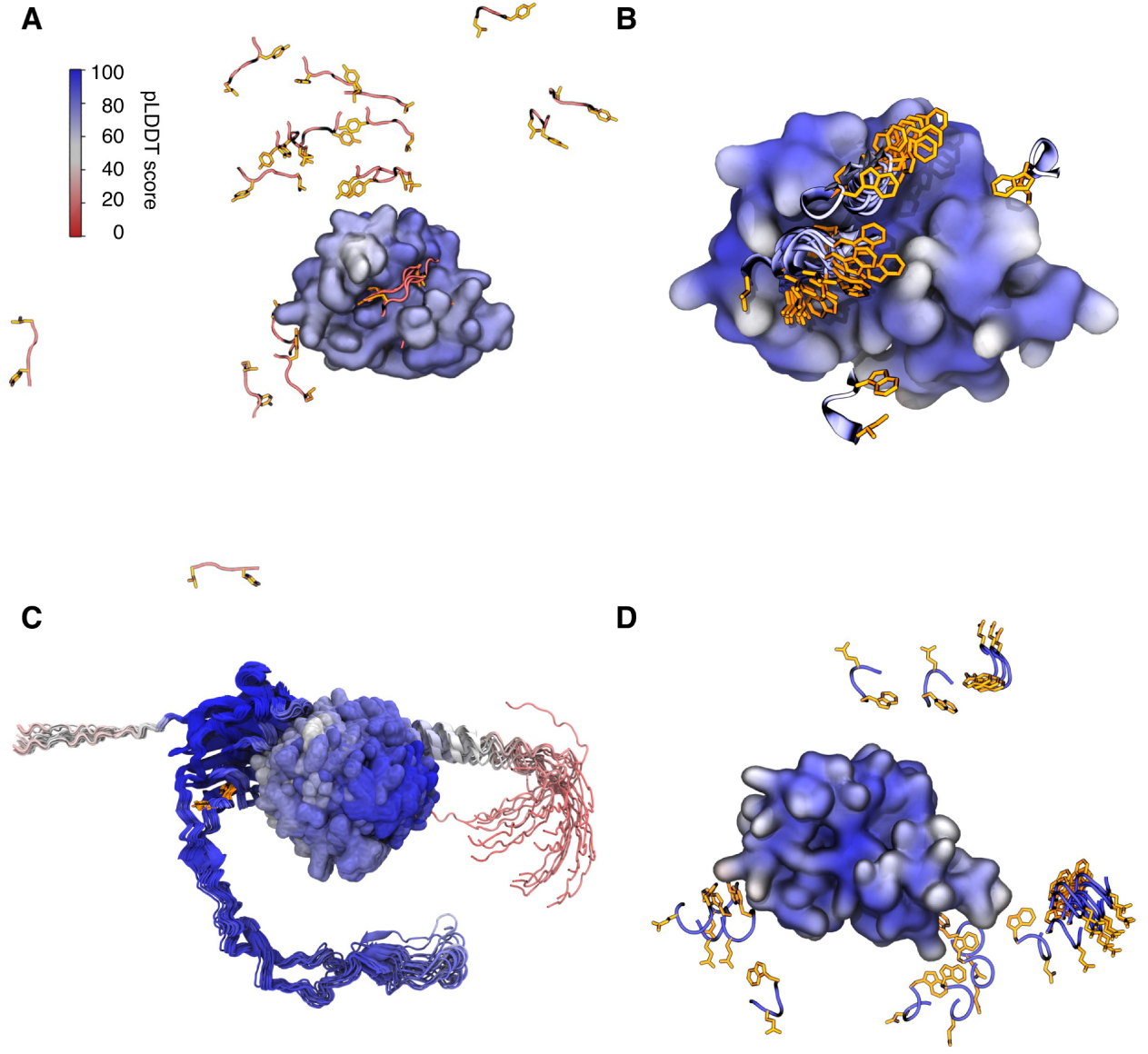
AlphaFold2.3 Multimer and AlphaFold3 increase the number of LC3-LIR interactions identified in predictions with full-length target proteins. (**A**) Atg8-Nup159 interaction is captured by AlphaFold2.3. Shown are 25 models of an AlphaFold2.3 prediction of the interaction between Atg8 (surface representation) and Nup159 (cartoon representation, with LIR residues highlighted as orange licorice), aligned on Atg8. Only the minimal LIR motif of Nup159 from the respective model, and only Atg8 from the top model are shown for clarity. AlphaFold2.2 did not predict this interaction in 25 models (data not shown). (**B**) ATG8E-Joka2 interaction is suggested by AlphaFold3. Shown are 25 models of an AlphaFold3 prediction of the interaction between ATG8E (surface representation) and Joka2 (cartoon representation, with LIR residues highlighted as orange licorice), structurally aligned on ATG8E. Only the minimal LIR motif of Joka from the respective model and only ATG8E from the top model are shown for clarity. AlphaFold2.2 and 2.3 did not predict this interaction in 25 models (data not shown). (**C**) AlphaFold2.3 does not capture GABARAP-calreticulin interaction. Shown are 25 models of an AlphaFold2.3 prediction of the interaction between GABARAP (surface representation) and calreticulin (cartoon representation, with LIR residues highlighted as orange licorice), aligned on calreticulin. (**D**) GABARAP-calreticulin interaction is not captured by AlphaFold3. Shown are 25 models of an AlphaFold3 prediction of the interaction between GABARAP (surface representation) and calreticulin (cartoon representation, with LIR residues highlighted as orange licorice), structurally aligned on GABARAP. Only the minimal LIR motif of calreticulin from the respective model, and only GABARAP from the top model are shown for clarity. All structures are colored by predicted local-distance difference test (pLDDT) score.

**Figure S2:**
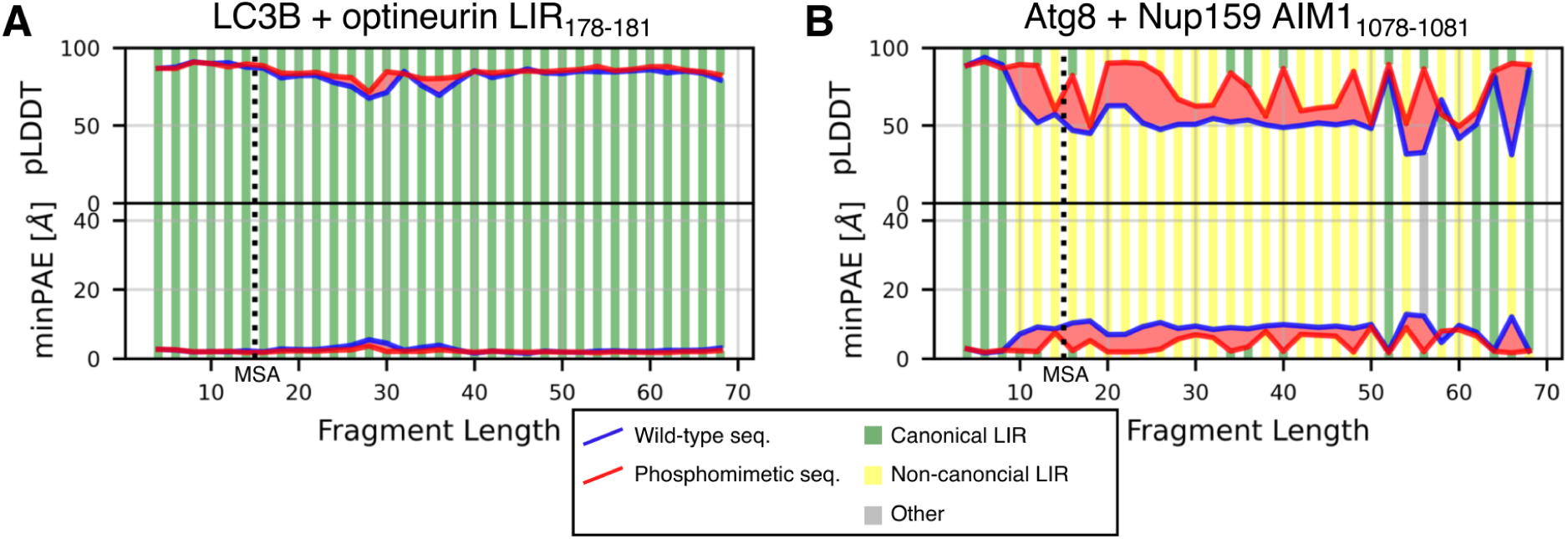
Fragment length and phosphomimetic mutations can modulate the prediction of LC3-LIR interactions with AlphaFold2. Predicted local-distance difference test (pLDDT) score, minimum predicted aligned error (minPAE), and binding mode for the four LIR/AIM residues in AlphaFold2.2 predictions of (**A**) LC3B and optineurin fragments containing the LIR_178_*_−_*_181_ and (**B**) Atg8 and Nup159 fragments containing the AIM1_1078_*_−_*_1081_. Fragments were varied (i) in length and (ii) by the introduction of phosphomimetic S/T to E mutations for experimentally confirmed phosphosites. The dotted lines indicate the minimum length required for the multiple sequence alignment (MSA) of AlphaFold2. Stripes above the pLDDT curve / below the minPAE curve indicate the binding mode of the respective fragment with phosphomimetic mutations (green: canonical; yellow: non-canonical; gray: other). Stripes below the pLDDT curve / above the minPAE curve indicate the binding mode of the respective wild-type fragment.

**Figure S3:**
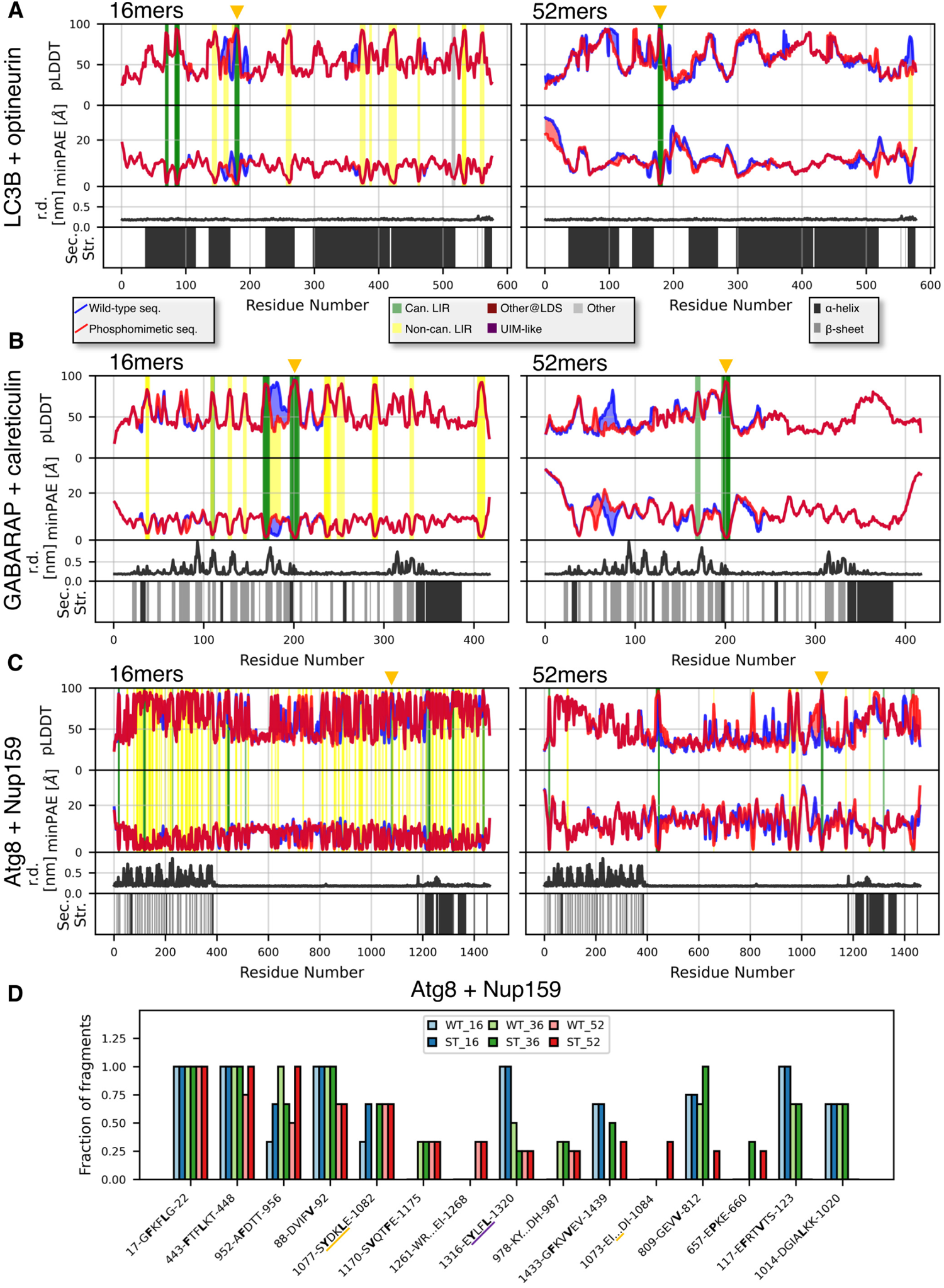
Fragment length modulates LC3-LIR complex predictions in the context of a systematic screen. Systematic screen of the interaction between 16- and 52-residue fragments of (**A**) optineurin screened against LC3B, (**B**) calreticulin screened against GABARAP, and (**C**) Nup159 against Atg8, respectively. The fragments have a 75% overlap. We performed the screen for the wild-type (WT) sequence, and a phosphomimetic (ST) sequence, in which S/T to E mutations were introduced at experimentally confirmed phosphosites. Predicted local-distance difference test (pLDDT) score, minimum predicted aligned error (minPAE), and binding mode for the interacting residues are shown (minimum minPAE from all fragments covering each residue and its respective pLDDT score). Stripes above the pLDDT curve / below the minPAE curve indicate the binding mode of the respective fragment with phosphomimetic mutations (see legend). Stripes below the pLDDT curve / above the minPAE curve indicate the binding mode of the respective wild-type fragment. Additionally, we calculated residue depth (r.d.) and secondary structure (sec. str.; for residues with pLDDT *>* 70) from the AlphaFold2-predicted structure for the full-length protein to add structural context to the sequence. The experimentally confirmed LIRs are indicated at the top by orange triangles. (**D**) Results of fragment-prediction screens for Nup159 combined with Atg8. Shown are the top fifteen predicted LIRs, sorted by relative occurrence in longer fragments (52mers *>* 36mers *>* 16mers). Residues binding in hydrophobic pocket 1 and 2 are highlighted in bold. The experimentally confirmed LIR is underlined in orange (found twice, in different binding modes), the experimentally non-functional LIR in purple.

**Figure S4:**
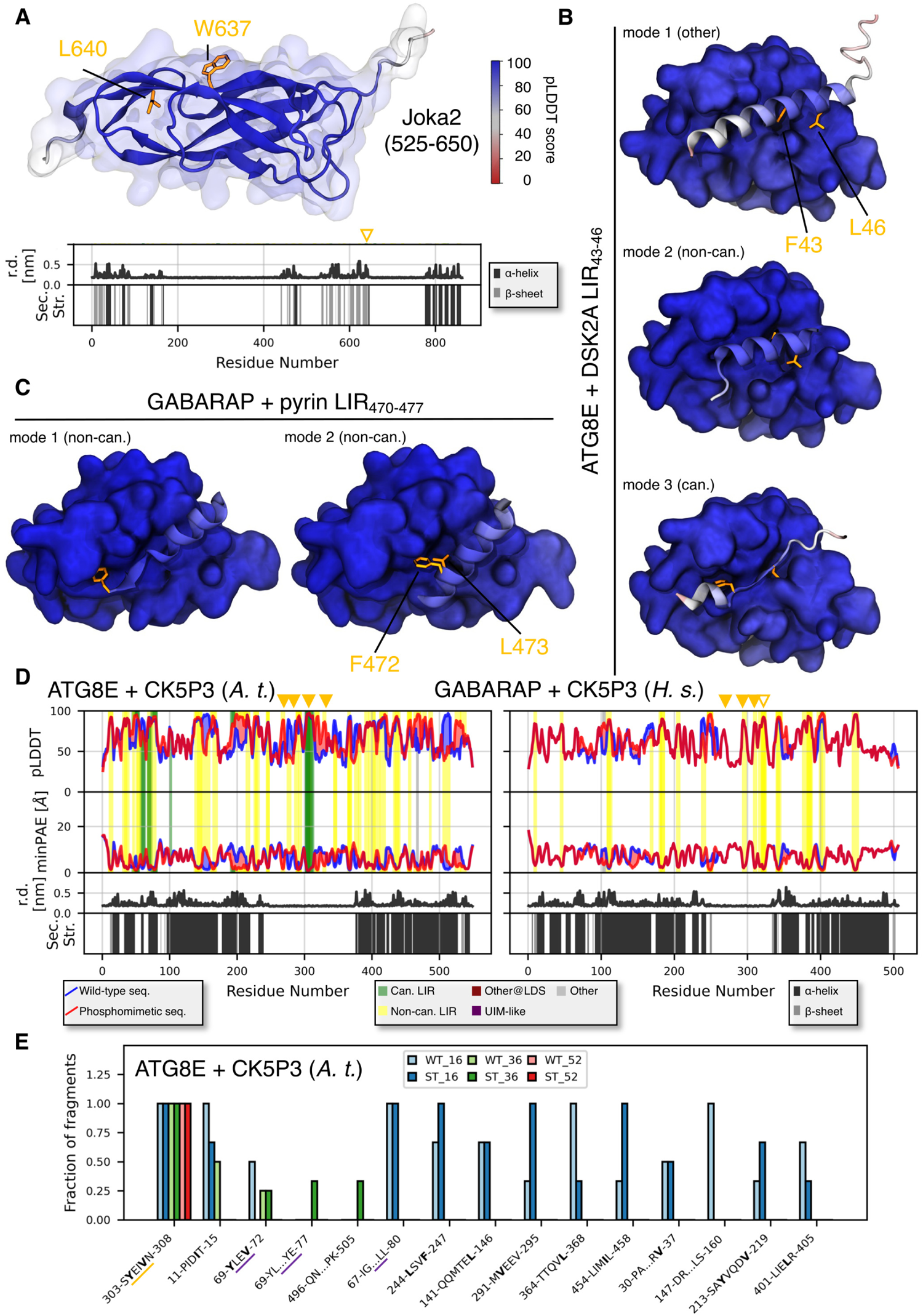
Detailed analysis of full-length and fragment structures predicted by AlphaFold2 Multimer assists in the interpretation of screen results. (**A**) Structural occlusion of the predicted LIR_637_*_−_*_640_ in Joka2 shown in the cartoon representation (W637 and L640 highlighted as orange licorice) of the full-length AlphaFold2 model (only the relevant domain is shown for visual clarity) and via calculated residue depth (r.d.) and secondary structure (sec. str.; for residues with pLDDT *>* 70). The predicted LIR_637_*_−_*_640_ is indicated via a hollow orange triangle. (**B**) Representative AlphaFold2 models of DSK2A fragments (cartoon representation) containing LIR_43_*_−_*_46_ (F43 and L46 highlighted as orange licorice) and interacting with ATG8E (surface representation) in different interaction modes. (**C**) Representative AlphaFold2 models of pyrin fragments (cartoon representation) (partially) containing LIR_470_*_−_*_477_ (F472 and L473 highlighted as orange licorice) and interacting with GABARAP (surface representation) in different interaction modes. Structures in **A**, **B**, and **C** are colored by predicted local-distance difference test (pLDDT) score.(**D**) Systematic screen of the interaction between 36-residue fragments of CK5P3 (*A. t.*) with ATG8E and CK5P3 (*H. s.*) with GABARAP, respectively. The fragments have a 75% overlap. We performed the screen for the wild-type (WT) sequence and a phosphomimetic (ST) sequence, in which S/T to E mutations were introduced at experimentally confirmed phosphosites. pLDDT score, minimum predicted aligned error (minPAE), and binding mode for the interacting residues are shown (minimum minPAE from all fragments covering each residue and its respective pLDDT score). Stripes above the pLDDT curve / below the minPAE curve indicate the binding mode of the respective fragment with phosphomimetic mutations. Stripes below the pLDDT curve / above the minPAE curve indicate the binding mode of the respective wild-type fragment. Additionally, we calculated residue depth (r.d.) and secondary structure (sec. str.; for residues with pLDDT *>* 70) from the AlphaFold2 predicted structure for the full-length protein to add structural context to the sequence. The experimentally confirmed LIRs are indicated at the top by filled orange triangles, the predicted LIR_320_*_−_*_324_ as a hollow orange triangle. (**E**) Results of fragment prediction screens for CK5P3 (*A. t.*) combined with ATG8E. Shown are the top fifteen predicted LIRs, sorted by relative occurrence in longer fragments (52mers *>* 36mers *>* 16mers). Residues binding in hydrophobic pocket 1 and 2 are highlighted in bold. The experimentally confirmed LIR is underlined in orange, the experimentally non-functional LIRs (in different binding modes) in purple.

**Figure S5:**
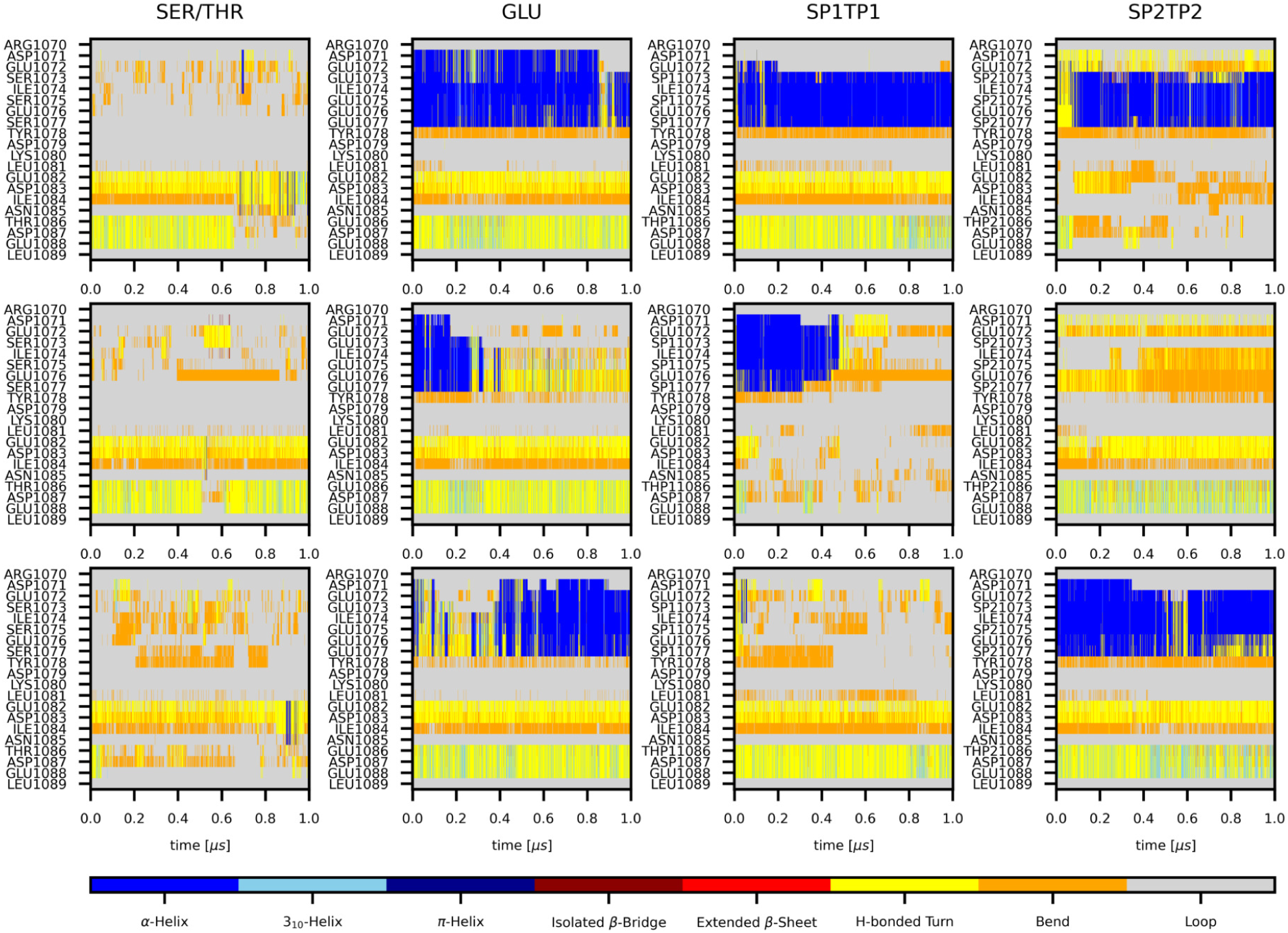
Nup159 AIM1 secondary structure in MD simulations. Plots show the secondary structure (colors as indicated in legend) of a 20-residue Nup159 fragment containing AIM1_1078_*_−_*_1081_ over time. Data from triplicate (1 µs each, rows 1-3) MD simulations of the respective fragment in complex with Atg8 for four different phosphorylation states (columns): unphosphorylated (SER/THR), phosphomimetic mutation (GLU), and phosphorylated (SP1/TP1: -HPO_4_*^−^* and SP2/TP2: -PO_4_^2^*^−^*, respectively).

**Figure S6:**
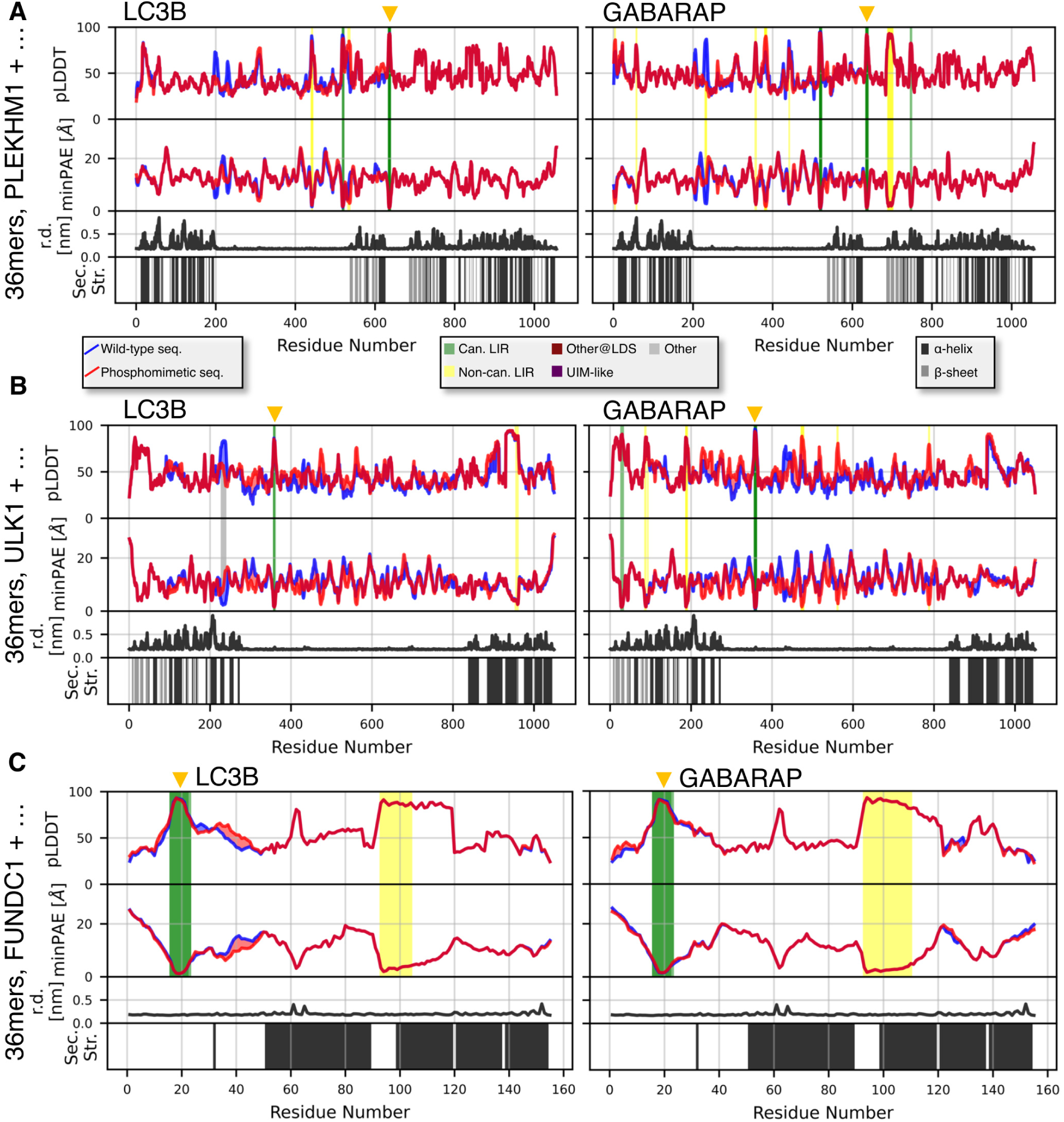
Sequence fragment scans for PLEKHM1, ULK1, and FUNDC1 with LC3B or GABARAP. Systematic screen of the interaction between 36-residue fragments of (**A**) PLEKHM1, (**B**) ULK1, and (**C**) FUNDC1 with LC3B and GABARAP, respectively. The fragments have 75% overlap with the adjacent fragments. We performed the screen for the wild-type (WT) sequence, and a phosphomimetic (ST) sequence, in which S/T to E mutations were introduced at experimentally confirmed phosphosites. Predicted local-distance difference test (pLDDT) score, minimum predicted aligned error (minPAE), and binding mode for the interacting residues are shown (minimum minPAE from all fragments covering each residue and its respective pLDDT score). Stripes above the pLDDT curve / below the minPAE curve indicate the binding mode of the respective fragment with phosphomimetic mutations. Stripes below the pLDDT curve / above the minPAE curve indicate the binding mode of the respective wild-type fragment. Additionally, we calculated residue depth (r.d.) and secondary structure (sec. str.; for residues with pLDDT *>* 70) from the AlphaFold2 predicted structure for the full-length protein to add structural context to the sequence. The experimentally confirmed LIRs are indicated at the top by orange triangles.

**Figure S7:**
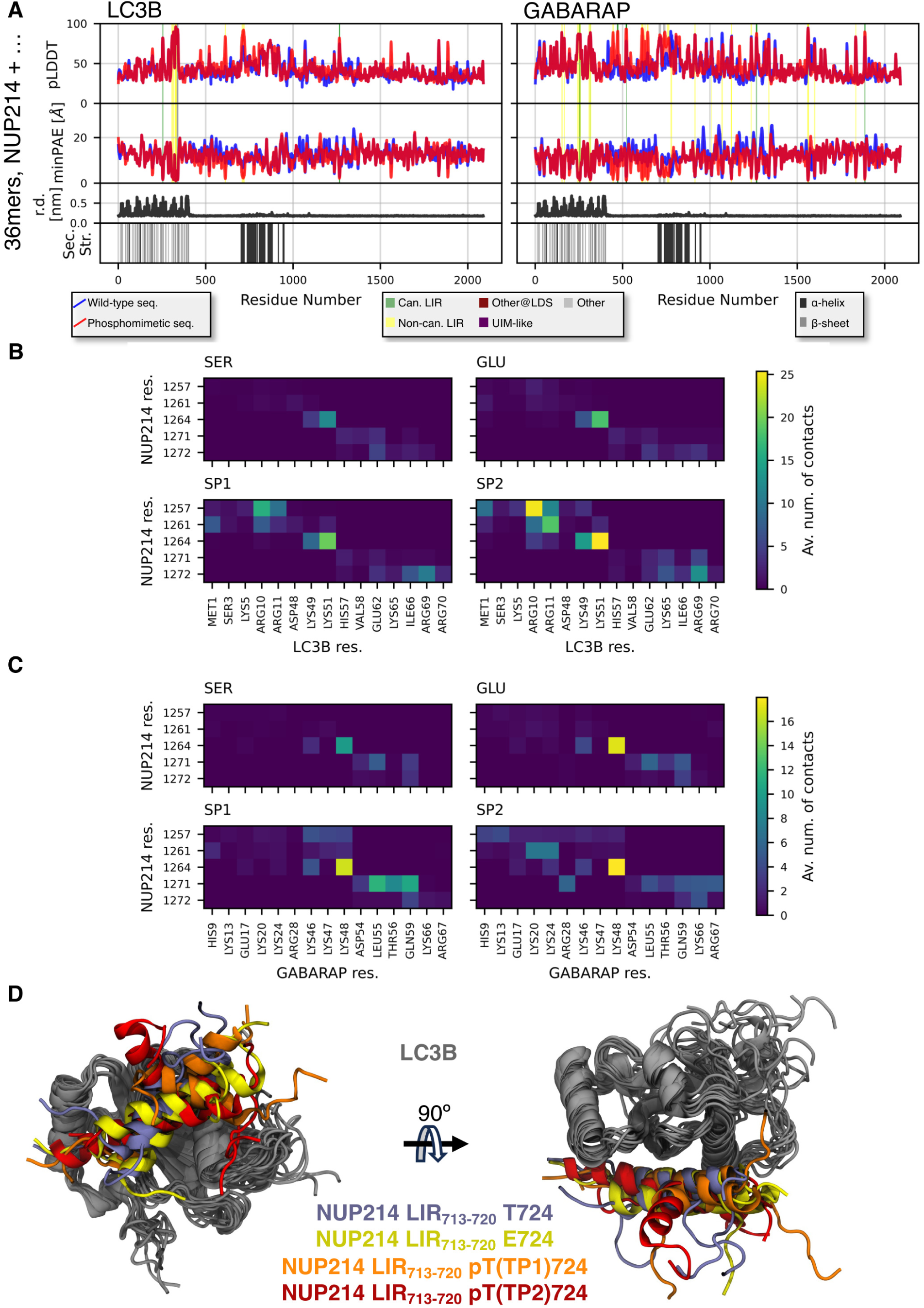
Phosphomimetic mutations and phosphorylations alter the interactions of predicted LIRs in NUP214 with LC3B and GABARAP. (**A**) Systematic screen of the interaction between 36-residue fragments of NUP214 with LC3B and GABARAP, respectively. The fragments have 75% overlap. We performed the screen for the wild-type (WT) sequence, and a phosphomimetic (ST) sequence, in which S/T to E mutations were introduced at experimentally confirmed phosphosites. Predicted local-distance difference test (pLDDT) score, minimum predicted aligned error (minPAE), and binding mode for the interacting residues are shown (minimum minPAE from all fragments covering each residue and its respective pLDDT score). Stripes above the pLDDT curve / below the minPAE curve indicate the binding mode of the respective fragment with phosphomimetic mutations. Stripes below the pLDDT curve / above the minPAE curve indicate the binding mode of the respective wild-type fragment. Additionally, we calculated residue depth (r.d.) and secondary structure (sec. str.; for residues with pLDDT *>* 70) from the AlphaFold2 predicted structure for the full-length protein to add structural context to the sequence. (**B and C**)Average number of heavy atom contacts (Av. num. of contacts; distance lower than 5 Å) per frame between potentially phosphorylated NUP214 residues and (**B**) LC3B and (**C**) GABARAP residues. Data from triplicate (1 µs each) MD simulations for four different phosphorylation states: unphosphorylated (SER), phosphomimetic mutation (GLU), and phosphorylated (SP1: -HPO_4_*^−^* and SP2: -PO_4_^2^*^−^*, respectively). (**D**) Final snapshots from 1 µs MD simulations of LC3B with a NUP214 fragment containing LIR_713_*_−_*_720_ in four different phosphorylation states (3 replicas per state): unphosphorylated (light blue), phosphomimetic mutation (yellow), and phosphorylated (TP1: -HPO_4_*^−^* and TP2: -PO_4_^2^*^−^*, orange and red respectively). All structures are aligned on LC3B. LC3B and the NUP214 fragment are shown in cartoon representation.

**Figure S8:**
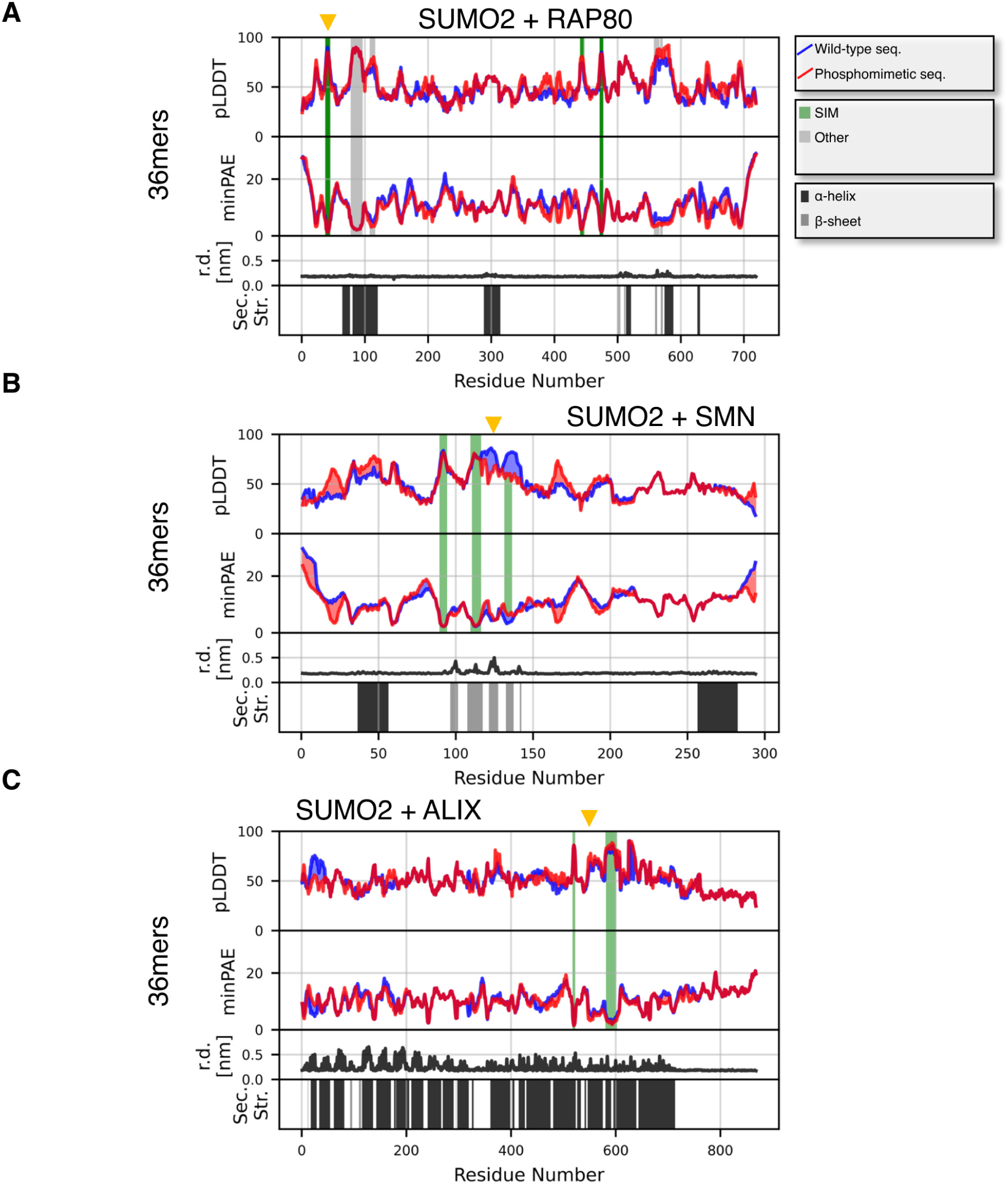
Sequence fragment scans for RAP80, SMN, and ALIX with SUMO2. Systematic screen of the interaction between 36-residue fragments of (**A**) RAP80, (**B**) SMN, and (**C**) ALIX with SUMO2. The fragments have 75% overlap. We performed the screen for the wild-type (WT) sequence, and a phosphomimetic (ST) sequence, in which S/T to E mutations were introduced at experimentally confirmed phosphosites. Predicted local-distance difference test (pLDDT) score, minimum predicted aligned error (minPAE), and binding mode for the interacting residues are shown (minimum minPAE from all fragments covering each residue and its respective pLDDT score). Stripes above the pLDDT curve / below the minPAE curve indicate the binding mode of the respective fragment with phosphomimetic mutations (see legend). Stripes below the pLDDT curve / above the minPAE curve indicate the binding mode of the respective wild-type fragment. Additionally, we calculated residue depth (r.d.) and secondary structure (sec. str.; for residues with pLDDT *>* 70) from the AlphaFold2 predicted structure for the full-length protein to add structural context to the sequence. The experimentally confirmed SIMs are indicated at the top by orange triangles.

**Table S1:**
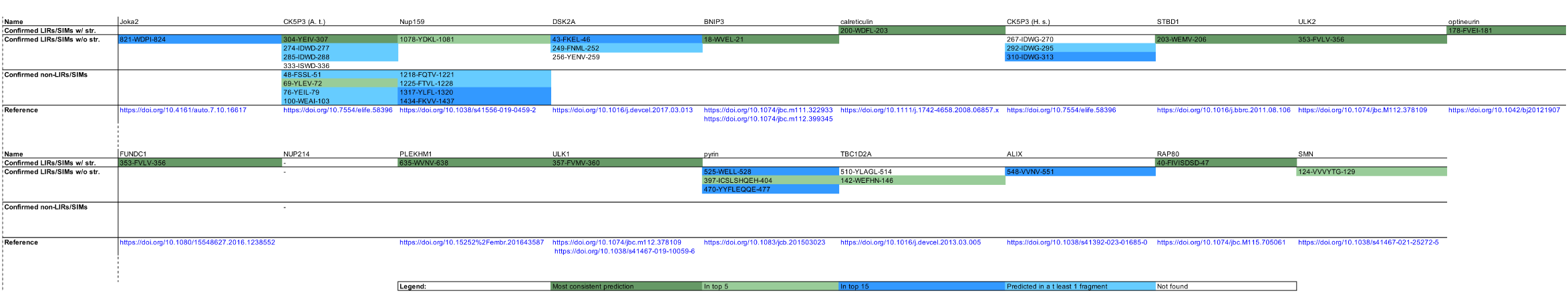
Overview over experimentally confirmed LIRs and SIMs for the proteins investigated with the fragment screen.

